# Search and Match across Spatial Omics Samples at Single-cell Resolution

**DOI:** 10.1101/2023.08.13.552987

**Authors:** Zefang Tang, Shuchen Luo, Hu Zeng, Jiahao Huang, Morgan Wu, Xiao Wang

**Affiliations:** Broad Institute of MIT and Harvard, Cambridge, MA, USA; Department of Chemistry, Massachusetts Institute of Technology, Cambridge, MA, USA

## Abstract

Spatial omics technologies characterize tissue molecular properties with spatial information, but integrating and comparing spatial data across different technologies and modalities is challenging. A comparative analysis tool that can search, match, and visualize both similarities and differences of molecular features in space across multiple samples is lacking. To address this, we introduce CAST (<u>C</u>ross-sample <u>A</u>lignment of <u>S</u>pa<u>T</u>ial omics), a deep graph neural network (GNN)-based method enabling spatial-to-spatial searching and matching at the single-cell level. CAST aligns tissues based on intrinsic similarities of spatial molecular features and reconstructs spatially resolved single-cell multi-omic profiles. CAST enables spatially resolved differential analysis (ΔAnalysis) to pinpoint and visualize disease-associated molecular pathways and cell-cell interactions, and single-cell relative translational efficiency (scRTE) profiling to reveal variations in translational control across cell types and regions. CAST serves as an integrative framework for seamless single-cell spatial data searching and matching across technologies, modalities, and disease conditions, analogous to BLAST in sequence alignment.

## Introduction

Spatial omics technologies enable direct profiling of gene expression and molecular cell types in intact tissues, organs^1–5^, and across different modalities like epigenomes^6^, translatomes^7^, and proteomes^8^. Computational approaches for spatial integration across technologies, disease conditions and modalities are essential for the analyses of spatial data, potentiating in-depth investigation of cell types, fates, and functional states in their spatial context. Analogous to sequence alignment in bioinformatics and atlas integration in single-cell omics, an ideal spatial integration tool for spatial omics should serve as a search engine and comparative analyzer to search, match, and visualize the similarity and differences among samples. Meanwhile, it should work robustly when dealing with vast numbers of cells, spanning various conditions and modalities. However, current spatial alignment methods^9^ can only handle small-scale, low-resolution, highly similar datasets using the same wet-lab technology, and fails to preserve accurate tissue morphology. On the other hand, image registration methods require landmark annotations and struggle with discrepancies in image properties. Moreover, effective fullstack spatial integration methods that allow accurate search-and-match of spatial omics data across technologies, modalities and disease conditions have not been achieved yet.

To address this, we introduce CAST (Cross-sample Alignment of SpaTial omics data), a spatial multi-modal integration approach for searching, matching, and visualizing the similarities and differences across spatial omics datasets. It leverages deep graph neural networks (GNNs) and physical alignment to harmonize spatial multi-omics data at the single-cell level while preserving cellular proximity in tissue niches. CAST can detect fine-grained common spatial features, perform robust physical alignment, and integrate samples of different spatial modalities, resolutions, and sizes. It is applicable across various low- and high-resolution spatial technologies (Visium, STARmap5, MERFISH2, RIBOmap7, Slide-seq3, Stereo-seq4), and can accurately match spatial samples of different sizes and gene numbers based on their inherent tissue properties, without supervision nor manual annotation of region of interest (ROI).

## Results

CAST is composed of three modules: CAST Mark, CAST Stack, and CAST Projection (Fig. 1a,b). First, CAST Mark leverages gene expression and spatial cell relationships to identify shared tissue architectures across samples using a deep GNN model. Second, CAST Stack performs high-resolution physical alignment of tissue samples based on the graph embedding of each cell, transforming the spatial coordinates of each tissue sample to a common tissue coordinate system. Third, using the well-aligned tissue coordinates, CAST Projection further enables single-cell resolved integration across the samples and even across different modalities of spatial omics.

**Fig. 1:**
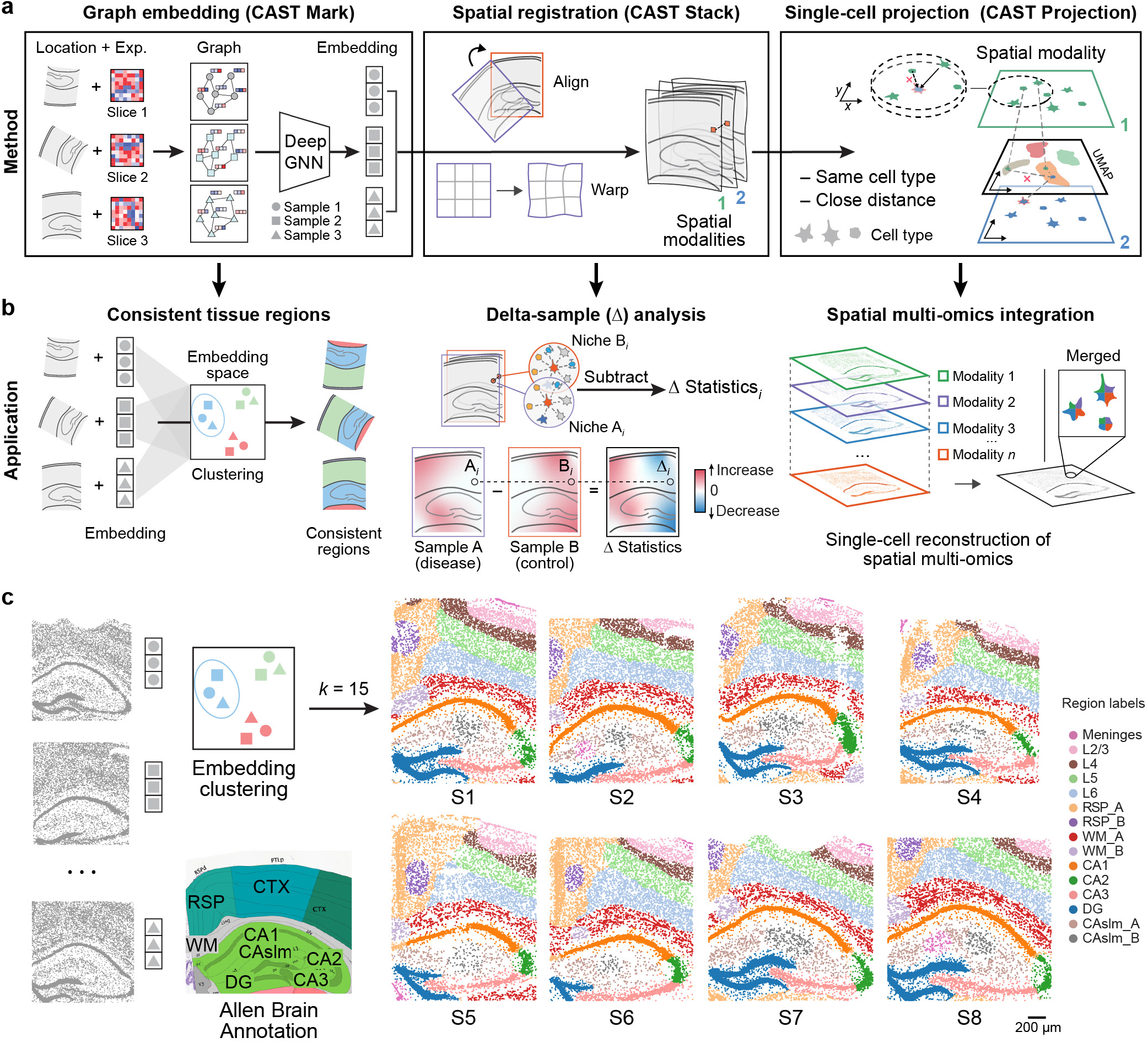
Schematic overview of CAST. **a**, The principles of the three modules in CAST. CAST Mark employs self-supervised graph learning on the coordinates and cell-by-feature matrix (such as single-cell gene expression) of the spatial dataset to obtain a shared spatial embedding across samples. CAST Stack aligns multiple different samples first by rigid registration then by free-form deformation (FFD) with the learned spatial features in CAST Mark. CAST Projection assigns single cells from the query samples into the reference sample by assigning single-cells from the query samples to the reference sample with the closest physical location and the most similar gene expression profile. **b**, The applications enabled by the three modules of CAST. CAST Mark identifies consistent regions across multiple samples by clustering the shared graph embedding. The consistent physical coordinates of multiple samples generated by CAST Stack enable cross-sample spatial analysis (named delta-sample analysis, ΔAnalysis) that detects the spatial differences of features (*e*.*g*. gene expression, cell-type compositions, cell-cell adjacencies, and cell-cell interactions) between conditions and visualize them as spatial gradients. Using physically-aligned, single-cell resolved cell assignment (named CAST Projection), CAST reconstructs a multi-omic tissue sample from multiple spatially resolved single-cell modalities. **c**, CAST Mark identifies consistent regions across multiple samples. The *k*-means (*k* = 15) clustering result of the graph embedding generated by CAST Mark across the samples S1–S8 in STARmap PLUS datasets (Supplementary Table 2), in comparison with Allen Brain Atlas17,18. Different colors in the cells indicate the different clusters of the graph embedding. CTX, Cerebral cortex; RSP, Retrosplenial area; WM, White matter; DG, Dentate gyrus; CAslm, CA stratum lacunosum-moleculare; CA1–3, hippocampal CA1–3 region; L2/3, L4, L5, L6, cortical layers 2/3, 4, 5 and 6, respectively.

### CAST Mark captures common spatial signatures across samples

Representing tissue samples using graphs^8^ shows the potential to overcome the inconsistent physical coordinates caused by different magnification, individual variation, and experimental batch effects. GNNs operate on graphs and have been recently used to learn representations of tissue organization of spatially resolved transcriptomics measurements^10–14^. However, traditional GNN architectures suffer from the over-smoothing problem that limits the depth of the network, raising doubts about their capability to capture large-scale continuities in tissue biology^14^. In addition, the traditional GNN architectures cannot identify the common spatial features across the samples in an unsupervised manner. To address these limitations, we created CAST Mark, a GNN model equipped with (1) GCNII layers which were designed to overcome the over-smoothing problem^15^, making the GNN learnable with a 9-layer depth; (2) a self-supervised learning objective (Extended Data Fig. 1a and Methods). By using the GCNII layers, CAST Mark overcomes the limited depth in a traditional GNN model and now has a large receptive field that enables unsupervised learning of spatial features using only single-cell gene expression profiles and physical cell coordinates as input, without the need for celltype or tissue-region annotations.

To evaluate the performance of the CAST Mark in learning the graph representations of cell locations across different samples, we first applied CAST Mark to a synthetic dataset consisting of one ground truth sample S1 from STARmap PLUS dataset^5^ and a simulated sample S1’ generated by applying random noise, feature dropouts, and global tissue distortion to sample S1 (Extended Data Fig. 1b and Methods). Each cell in the simulated sample S1’ has a one-to-one ground-truth partner cell in sample S1. We performed *k*-means clustering on the graph embedding to examine whether CAST Mark could retain the shared spatial information between S1 and S1’. Although the graph structures of S1 and S1’ are different due to added random noise, the regional patterns are consistent across samples in both the physical space (Extended Data Fig. 1c) and the graph embedding space (Extended Data Fig. 1d). These observations are confirmed by quantitative analysis, where 20 clusters show a high adjusted rand index (ARI = 0.85), and 92% of cells in S1’ belong to the same clusters as its ground-truth partners in S1 (Extended Data Fig. 1e). Notably, even when increasing the number of clusters *k* to 100, the clustering results still show a considerable cross-sample consistency both by visual inspection and quantification (Extended Data Fig. 1e,f; ARI = 0.43, consistent cell percentage = 59%). Furthermore, despite different clustering parameters (10–100), each cell is still physically adjacent (average distance = 6.95 μm, smaller than the typical size of a cell) to the correct clusters (Extended Data Fig. 1g), suggesting the robust performance of CAST Mark despite sample variability.

To demonstrate the technical advances of the CAST Mark GNN, we compared the performance of CAST Mark with published GNN-based tissue segmentation methods, SpaGCN^11^, STAGATE^12^, and GraphST^16^. Notably, none of these GNNs can directly generate consistent graph embeddings across multiple samples, so we benchmarked the GNN performance on a single sample (Extended Data Fig. 1h-i and Supplementary Table 1). While CAST Mark outperforms existing methods in terms of resolution and contiguity in sample S1 (Extended Data Fig. 1h), it furthermore scales up to a mouse half brain coronal sample containing ∼60,000 cells (Extended Data Fig. 1i).

Encouraged by the cross-sample consistency of CAST Mark graph embedding trained on the synthetic dataset, we next examined whether CAST Mark could achieve consistent label-free segmentation with real biological samples. We applied CAST Mark to the 2,766-gene STARmap PLUS dataset^5^ composed of 8 coronal brain slices near the hippocampus region (named as slice S1–S8) from multiple mice with different conditions, ages and strains (Supplementary Table 2). *K*-means clustering (*k* = 15) yielded consistent tissue-region identification across 8 samples (Fig. 1c), which agreed well with existing knowledge of mouse brain anatomy^17,18^. Notably, the CAST Mark region segmentation result trained on 8 biological samples is better than that trained on a single sample S1 (Extended Data Fig. 1h), suggesting the ability of CAST Mark to gain performance by summarizing real biological variations shared by all samples and overcoming individual variations unique to each sample. We further tested an extremely high clustering resolution by 100-class *k*-means clustering (*k* = 100), and the results still showed remarkable consistency across the 8 samples (Extended Data Fig. 2a), suggesting the intrinsic ability and power of CAST Mark learning scheme in resolving fine tissue architectures consistent across all samples, although the biological meaning of those fine clusters warrant further investigation. These results confirm that unsupervised identification of anatomical tissue regions is achievable solely using spatially resolved molecular features.

Finally, to investigate whether the identified tissue regions share similar characteristics across samples, we first compared the cell-type compositions of each canonical region (*k*-means cluster, *k* = 15), which were highly consistent across the 8 samples (Extended Data Fig. 2b). Next, we investigated the specific genes expressed in each region (Extended Data Fig. 2c and Methods). These regional marker genes, such as *Cux2* in cortical layers 2/3 (L2/3), *Tshz2* in retrosplenial cortex (RSP), *Prox1* in dentate gyrus (DG) region, and *Tenm3* in the hippocampal CA1 region (CA1), share similar spatial expression patterns across samples, which is consistent with the Allen ISH database^17^ (Extended Data Fig. 2c,d and Supplementary Table 3). Notably, the consistent patterns of gene expression and cell-type abundance across the 8 samples strongly support that CAST can robustly identify the concordant and biologically meaningful spatial features across different samples with biological and individual variations, which are further used as a foundation for sample alignment.

### CAST Stack performs robust physical alignment across samples

We next developed the CAST Stack module to automatically align and overlay two different samples into a consistent physical coordinate system. As the cytoarchitecture of tissue samples falls on a spectrum between completely stereotypical to random, an ideal alignment method should meet the following requirements: (1) robust correction of local differences in batches, conditions, tissue morphology, and experimental technologies; (2) preservation of cellular organization inside the tissue.

Since CAST Mark is capable of generating common graph embeddings for cells across multiple samples, we hypothesize that the similarity of cellular graph embeddings reflects the physical proximity of the cells in tissues, and thus can be used to physically register one query tissue sample to the reference sample. To test this, we used the synthetic sample (S1’) as the query and the ground truth sample (S1) as the reference. Given one cell in the query sample, we calculated the Pearson correlation (*r*) between the graph embeddings of the query cell and all the cells in the reference sample. We found that ground truth pairs between S1 and S1’ show a strong correlation (average *r* = 0.97; Fig. 2a), while randomly chosen cell pairs show little correlation (average *r* = 0.04; Fig. 2a). When plotted in the physical space, cells in the reference sample that are highly correlated with the query cell are predominantly localized around the ground truth reference cell, especially within the same tissue region (Fig. 2b and Extended Data Fig. 3a).

**Fig. 2:**
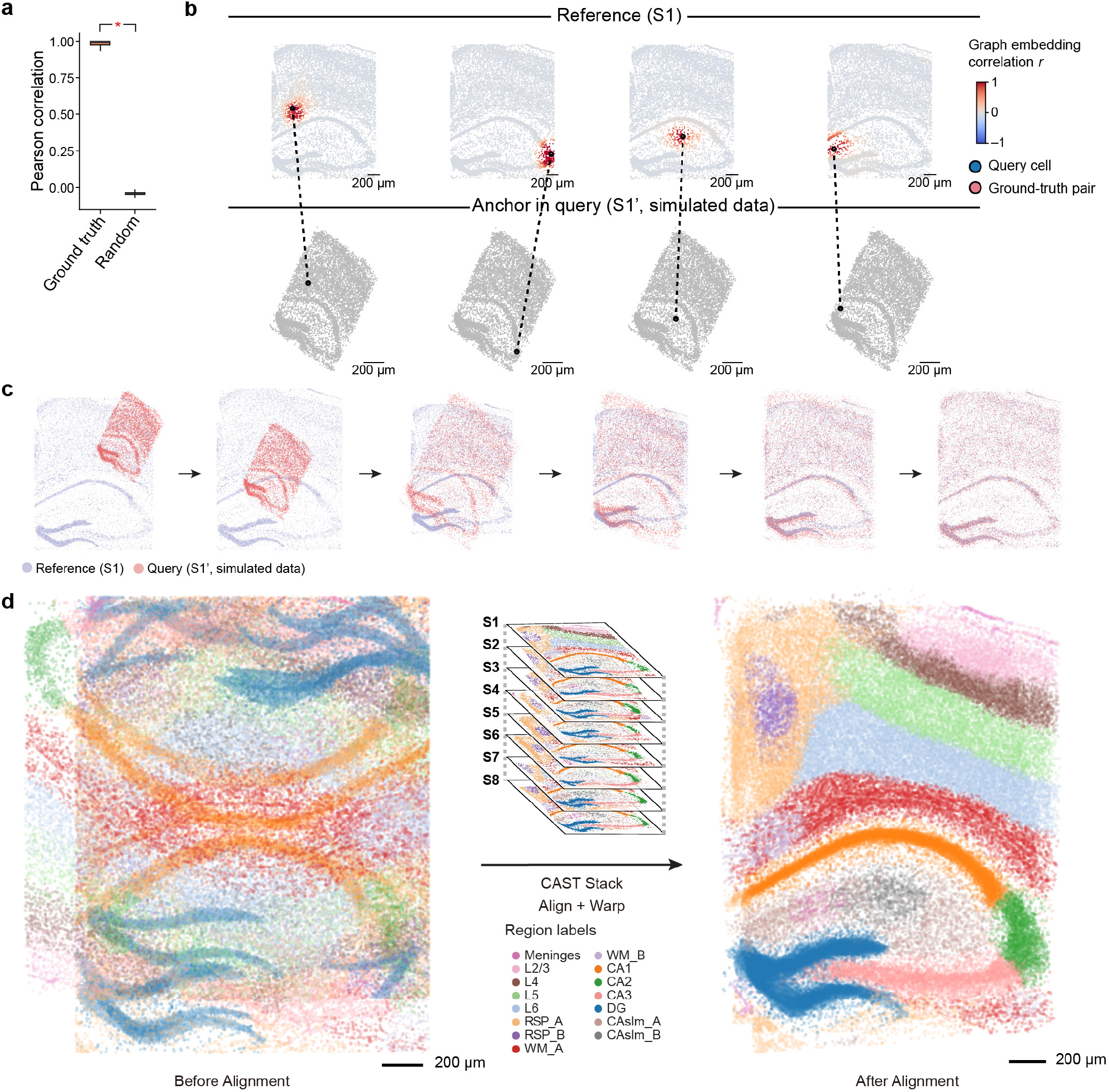
CAST Stack automatically aligns tissue samples from biological replicates. **a**, The boxplot shows a significantly higher (One-way ANOVA) Pearson correlation of graph embedding of the ground truth cell pair (Ground-truth, mean = 0.97, *n* = 8,789) than the ones in random distribution (Random, mean = 0.04, *n* = 8,789) between samples S1’ and S1. **b**, Given the query cell in the query sample (simulated dataset S1’), the cells in the reference sample (S1) are colored by Pearson correlation of the graph embedding between the reference cells and the given query cell. **c**, Schematic demonstration of the CAST Stack alignment process. The alignment process of the simulated dataset S1’ (query sample) and S1 (reference sample) is visualized. **d**, CAST Stack aligns 8 samples (S1–S8) into a consistent physical coordinate system. Cells are colored by tissue region labels generated by CAST Mark, the same as is shown in Figure 1c.

Based on this observation, we concluded that the cross-sample correlations of cell pairs could predict their probable match of tissue locations. However, due to the inherent anatomical diversity across samples, we would lose the cell organization if we simply assigned each query cell to the position with the highest similarity of the graph embedding. Therefore, we designed a gradient descent (GD)-based approach to minimize overall cell location differences while preserving tissue structure during alignment transformations, by maximizing the sum of similarity between each query cell and its nearest reference cell (Methods). Instead of building alignment by satisfying every cell at its optimum, CAST Stack prioritizes preserving biologically meaningful tissue structure and avoids local minimums possibly derived from stochastic sample variations. We designed the CAST Stack alignment as a two-phase process. During the first phase, only global affine transformation is allowed. After affine transformation roughly aligns the samples, in the second phase, CAST Stack utilizes free-form deformation (FFD), a powerful constrained non-linear warping approach, to handle local morphological differences among tissue samples.

We then applied this soft registration strategy to the S1’-S1 query-reference pair (Fig. 2c). Despite large structural and shape differences introduced in S1’, the two samples were accurately aligned according to the high spatial correlations (Pearson *r* of graph embeddings between cells, same below unless otherwise stated) between the query cells and their nearest neighbors in the reference slice (Extended Data Fig. 3b). After the soft registration, the physical distances between the ground truth pairs (average distance = 38 μm; Extended Data Fig. 3c) are significantly smaller than the random pairs (average distance = 1,133 μm; Extended Data Fig. 3c), confirming that CAST can precisely align two different slices into consistent tissue coordinates.

Next, we applied CAST Stack to the 8 hippocampal brain samples (S1–S8) from different mice with varied tissue morphologies, ages, and conditions^5^. We selected S1 as the reference slice, and subsequently aligned S2–S8 to S1 using CAST Stack. Similar to the S1’-S1 query-reference pair, cells from S2–S8 have the highest spatial correlation with cells from S1 at the corresponding tissue locations, especially within the same cluster of graph embeddings from CAST Mark (Extended Data Fig. 3d). After alignment, all the cells in the query samples (S2–S8) are transformed to the same physical coordinate system defined by the S1 reference (Fig. 2d and Extended Data Fig. 3e). The high correlation between the query cells with its closest physical neighbor cell in the S1 reference (Extended Data Fig. 3e) suggests CAST Stack properly aligns each sample through soft registration while preserving the cellular organization of the tissues.

To demonstrate the wide utility of CAST, we applied CAST on different spatial technologies, such as Visium, Stereo-seq, MERFISH and Slide-seq. Samples with similar size can be efficiently aligned not just within a single technology but also across multiple different technologies (Fig. 3a and Extended Data Fig. 4a-d; Supplementary Table 4). Remarkably, alignment of three different technologies can be achieved effortlessly in a single run (Fig. 3a). Moreover, when utilizing CAST to align a small slice (hippocampus and partial cortical region) with larger half-brain slices measured by different spatial technologies and size (STARmap, MERFISH and Slide-seq), we found CAST can independently search and match the small slice to the large one without manually specifying region of interest (ROI) nor annotating of landmarks (Fig. 3b-d and Supplementary Video 1). Additionally, we also tested the performance of CAST Mark and CAST Stack with limited gene panels. CAST can successfully align two STARmap samples collected with a 64-gene panel (S64_1 and S64_2; Extended Data Fig. 4e). CAST can also align samples with drastically different gene panels with only a few overlapping genes, showcased by the successful alignment of a 64-gene sample to a 2766-gene sample (S64_1 and S1; Fig. 3e).

**Fig. 3:**
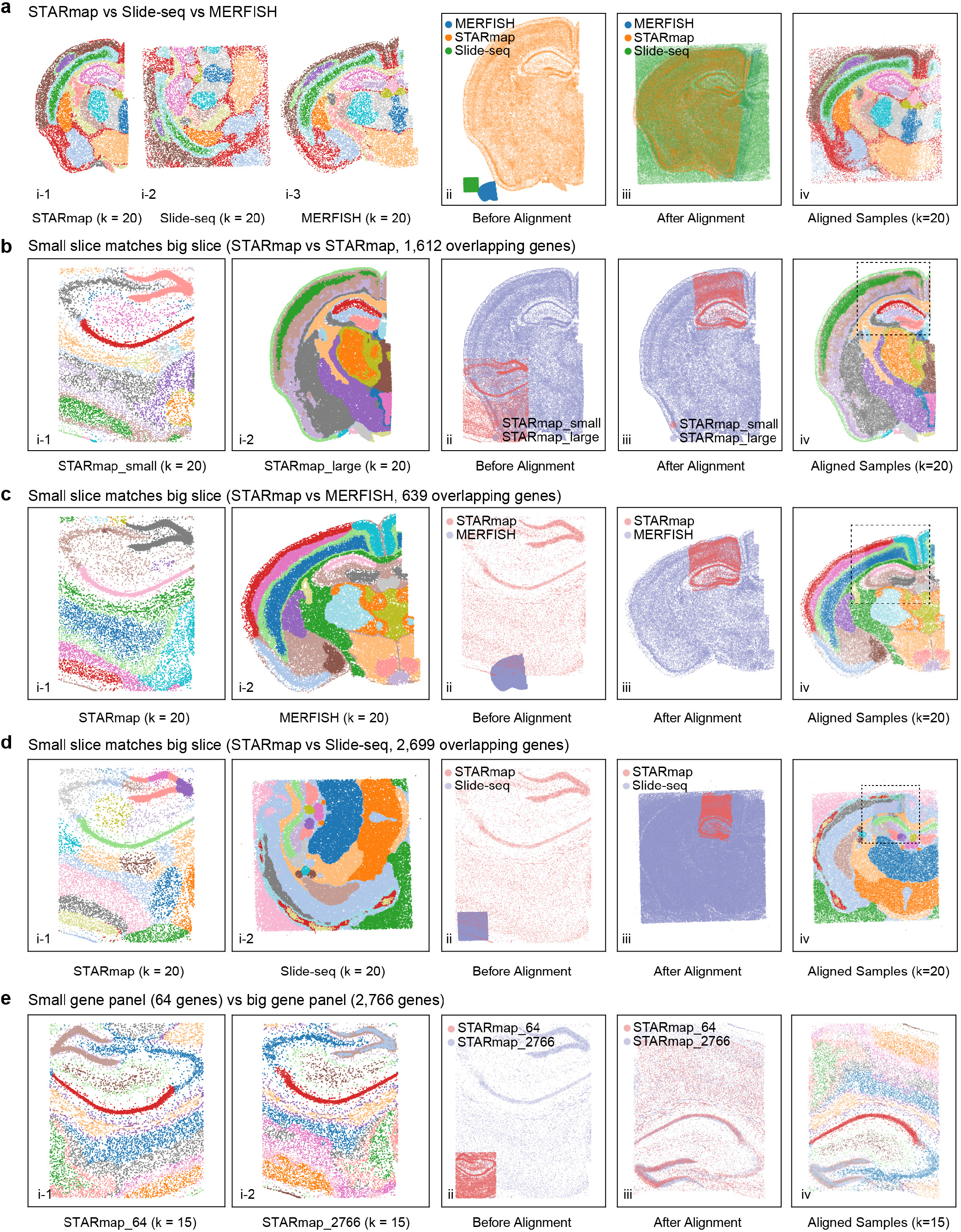
CAST aligns tissue samples across spatial technologies regardless of different tissue areas and gene panel sizes. **a**, CAST integrates three samples of Slide-seq, MERFISH, and STARmap (STARmap_mouse17), respectively, generating a common physical coordinate system. Panels **i**-**iv** indicate: **i**, the joint *k*-Means clustering results of CAST Mark graph embeddings of different samples, colored by joint clusters. **ii-iii**, spatial coordinates of the samples before (**ii**) and after (**iii**) alignment. **iv**, aligned samples colored by joint *k*-Means clustering results of the graph embeddings. **b-d**, CAST automatically searches and matches small slice to big slice across technologies at high spatial resolution (**b**, STARmap vs. STARmap. STARmap_small is S1 and STARmap_large is STARmap_mouse1. **c**, STARmap (S1) vs. MER-FISH; **d**, STARmap (S1) vs. Slide-seq). The panel order is analogous to **a. e**, CAST Stack aligns the samples with different gene panels designed by different probes. STARmap_64 is S64_1 (Supplementary Table 2) and STARmap_2766 is S1. Two samples share only 64 genes. The panel order is the same as **a**.

Furthermore, we compared CAST with the existing spatial alignment tool PASTE, which adopts optimal transport (OT) to perform only global affine transformation to align voxel-based spatial transcriptomics data^9^. The results showed that PASTE failed to align S2-8 with S1 in both pairwise alignment (Extended Data Fig. 5a) and multi-slice alignment (Extended Data Fig. 5b) tasks, even after human-supervised flipping and rotation (Extended Data Fig. 5c). Meanwhile, it failed to align the spatial datasets with a large number of the cells or voxels (Supplementary Table 5).

### Delta-sample (Δ) analysis reveals the spatially heterogeneous disease features

Traditional single-cell analysis workflows can be adapted to find significant differences between samples, such as celltype abundance, differential gene expression, and cell-cell interactions in the spatial transcriptomics data^5^. However, by preserving single-cell resolved spatial relationships, it is possible to interrogate the continuous spatial gradients of such differences in cellular neighborhoods across multiple samples with unified tissue coordinates^19^ (Fig. 4a). Here, enabled by the physical alignment of CAST Stack, we further introduce a new spatial omics analysis strategy, delta-sample analysis (ΔAnalysis; Methods), to uncover comparative spatial heterogeneity across tissue samples: (i) given a cell and a physical radius (*R*), we first define a cell-centered neighborhood termed spatial niche; (ii) we then analyze the local difference of interrogated features between samples within *R*, such as cell abundance (ΔCell), gene expression (ΔExp), cell-cell adjacency (ΔCCA), and cell-cell interaction (ΔCCI, e.g. ligand-receptor interactions), which can be visualized as spatial gradient maps (Fig. 4a); (iii) by aggregating the local Δ features of single-cells throughout the replicates and samples, we conduct statistical analysis at single-cell level to test whether there is a significant difference of spatially resolved features between samples.

**Fig. 4:**
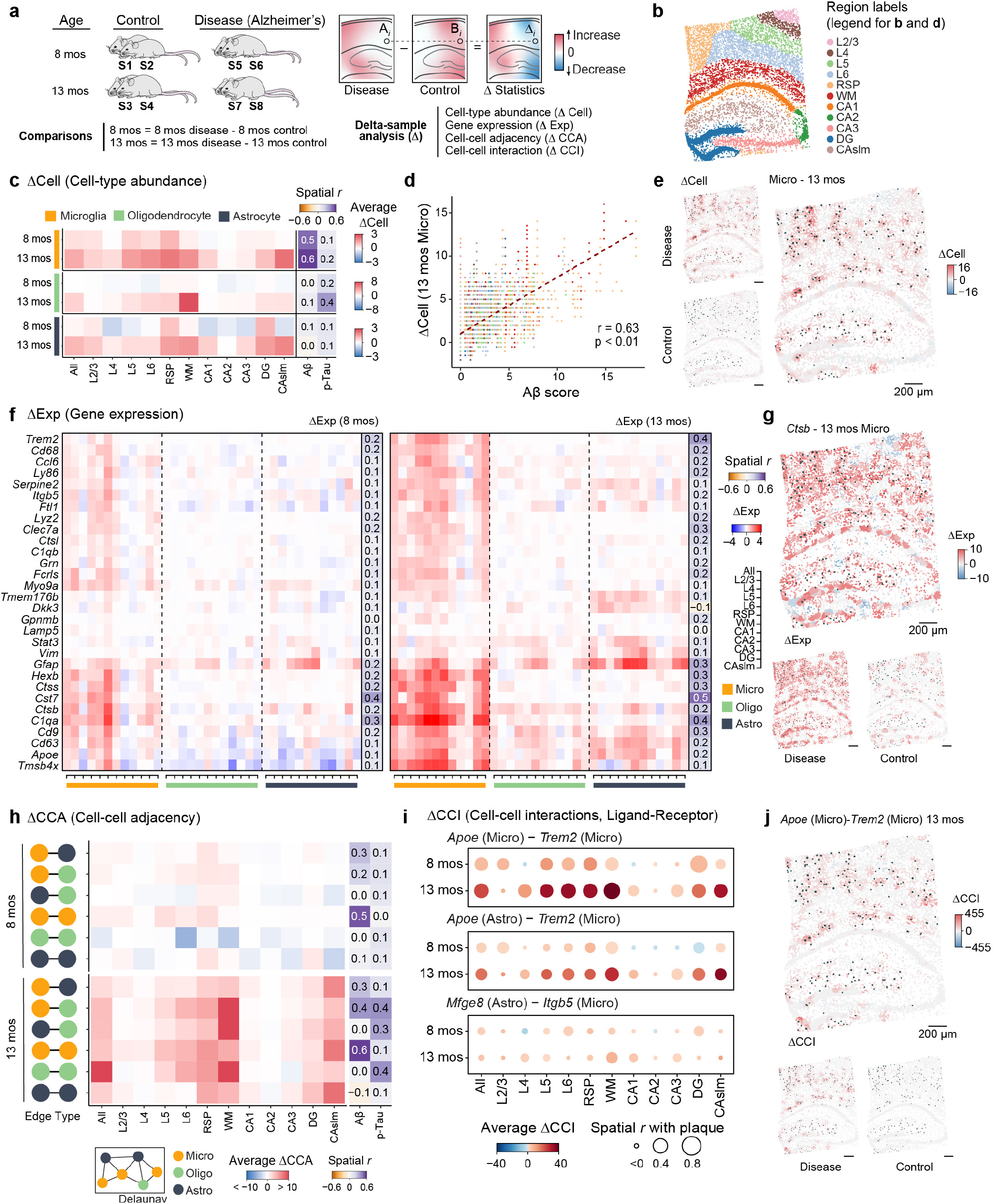
Delta-sample analysis (ΔAnalysis) detects spatial differences of molecular characteristics between disease and normal conditions. **a**, ΔAnalysis is performed to interrogate the spatial differences of cell type abundance (ΔCell), gene expression (ΔExp), cell-cell adjacency (ΔCCA), and cell-cell interaction (ΔCCI), between different conditions. Two comparisons used for analysis (Supplementary Table 2): 8 mos, S5, S6 (disease samples) and S1, S2 (control samples) in the 8-month group; 13 mos, S7, S8 (disease samples) and S3, S4 (control samples) in the 13-month group. The cell-centered neighborhood termed spatial niche is defined given a cell and a physical radius (*R*). The local difference of interrogated features (Δ statistics) can be calculated by the subtraction of the values in the niches with the same location but different samples (Niche Ai in the disease sample and Niche Bi with the same location in the control sample). The difference can be visualized as spatial gradient maps. **b**, The niche centers (cell) colored by the different regions. **c**, The average delta cell type abundance (ΔCell) in different regions, cell types and comparisons. The average spatial correlation (Pearson *r*) between the ΔCell and the Aβ-plaque as well as p-Tau scores (Methods) are displayed aside. For visualization, The values of 4 combinations in each comparison are averaged (13 mos comparison: S7 − S3, S7 − S4, S8 − S3, S8 − S4; 8 mos comparison: S5 − S1, S5 − S2, S6 − S1, S6 − S2). **d**, The scatter plot shows the significant high spatial correlation between the ΔCell (Microglia) and the Aβ-plaque score in S8 sample (Pearson correlation test, *r* = 0.63, n = 10,372). The ΔCell values (*y* axis) are the subtraction of the cell type abundance of S8 and the average values of the S3 and S4 (S8 − (S3 + S4) / 2). **e**, The spatial gradient map (S8 coordinates) shows the ΔCell pattern of the microglia in 13 mos comparison (S8 − (S3 + S4) / 2). The disease sample shows the microglia abundance in S8, while the control one (plotted in the S8 coordinates) shows the average values of the S3 and S4. The dark green dots represent the Aβ-plaque in the S8 sample, and the size of the dots indicates the area of the Aβ-plaque. **f**, The average delta gene expression (ΔExp) of the plaque-induced genes in different regions, cell types and comparisons. The average spatial correlation (Pearson *r*) between the overall (gene expression in all cells) ΔExp and the Aβ-plaque score are listed aside. Analogous to **c**, the values of 4 combinations in each comparison are averaged. Micro, Microglia; Oligo, Oligodendrocyte; Astro, Astrocyte. **g**, Analogous to **e**, the ΔExp of the *Ctsb* gene in microglia and 13 mos comparison (S8 − (S3 + S4) / 2). **h**, Analogous to **c**, the delta cell-cell adjacency (ΔCCA) pattern. **i**, Analogous to **c**, the delta cell-cell interaction (ΔCCI) pattern of the *Apoe* (Micro) - *Trem2* (Micro), *Apoe* (Astro) - *Trem2* (Micro) and *Mfge8* (Astro) - *Itgb5* (Micro) in different regions and comparisons. **j**, Analogous to **e**, the ΔCCI pattern of the *Apoe* (Micro) - *Trem2* (Micro) in 13 mos comparison (S8 − (S3 + S4) / 2).

To demonstrate ΔAnalysis, we analyzed the STARmap PLUS data from Alzheimer’s disease (AD) brain samples with spatially resolved single-cell RNA expression of 2,766 genes and protein histopathological staining of amyloid beta (Aβ) plaques and hyperphosphorylated tau (p-Tau) tangles across 4 diseased samples (TauPS2APP mouse model) and 4 control samples (age-matched wild-type mouse)^5^. The 8 samples are grouped into two comparisons: 8-month disease versus control (8-mos) and 13-month disease versus control (13-mos; Fig. 4a).

First, we interrogated the differences in cell-type abundances within niches (ΔCell) between diseased and control samples. When constructing niches, we assigned each cell-centered niche (*R* is set as 50 μm) with a region label based on the CAST Mark (Fig. 4b). We then calculated ΔCell in the aligned cross-sample niches for three disease-relevant glial cell types: astrocytes, microglia, and oligodendrocytes. The ΔCell of oligodendrocyte shows the most significant changes in the white matter (WM) region in the 13-mos group and correlates with the accumulation of p-Tau intensity (Fig. 4c and Extended Data Fig. 6a), consistent with the previously reported correlation between the oligodendrocytes and the p-Tau^5,20^. The abundance of microglia increased in retrosplenial cortex (RSP), cortical layer 5 (L5), and cortical layer 6 (L6) regions at 8 months, and kept increasing in all regions except the hippocampal CA2 and CA3 region (CA2 and CA3) at 13 months (Fig. 4c). The elevated abundance of the microglia in disease versus control quantified by ΔCell shows a high spatial correlation with the density of the increased Aβ plaques (average Pearson *r* = 0.47 in 8-mos; average Pearson *r* = 0.55 in 13-mos; Fig. 4c-e and Extended Data Fig. 6b,c), further validating the activated response of the microglia to the Aβ plaque^21,22^. The ΔCell values with different radii (*R* from 5 μm to 200 μm) show a consistent spatial pattern (Extended Data Fig. 6d).

Next, we calculated the delta-sample single-cell gene expression changes (ΔExp) in each niche to interrogate the spatial gradient of differential gene expression. Further correlating ΔExp with spatial locations of the Aβ-plaque enables us to identify spatially resolved plaque-induced genes (PIGs; Methods). Based on this strategy, we identified 30 spatially resolved PIGs (15 PIGs in the 8-mos group; 27 PIGs in the 13-mos group; Fig. 4f and Extended Data Fig. 6e; Supplementary Table 6). These genes are over-expressed in the disease samples, and their ΔExp patterns are spatially correlated with Aβ plaques (Extended Data Fig. 6f) and enriched in peptide binding (GO:0042277) and lysosome pathway (mmu04142; Extended Data Fig. 6g; Supplementary Table 6). Notably, our results are consistent with previously reported PIGs identified by Spatial Transcriptomics (ST) technology^21^ and the initial STARmap PLUS study (Fisher Exact test, p < 0.001), such as *Apoe, C1qb, Cd63, Ctsb*, and *Gfap*. More importantly, the ΔExp analysis further uncovered the regional heterogeneity of changes in PIG expression levels in AD. For instance, *Ctsb*, an AD-related gene^23^, does not show uniform over-expression in the microglia across the sample, rather it exhibits over-expression in the CA stratum lacunosum-moleculare (CAslm), L6, and RSP regions in the 13-mos comparison (Fig. 4f,g; Extended Data Fig. 6h).

To investigate the disease-associated cell-cell interactions, we first examined differential cell-cell adjacency^24^ (ΔCCA; Fig. 4h) defined by the edges of the cellular graph. The ΔCCA of the oligodendrocytes with other glial cells increased remarkably in the white matter and exhibited high spatial correlation with p-Tau, hinting that the increased p-Tau may activate the interactions among the glial cells. Meanwhile, the varied adjacency patterns of the microglia with other glial cells also showed a high spatial correlation with Aβ-plaque. These disease-associated CCA patterns encouraged us to further investigate the cell-cell interactions^25^ among the glial cells. To this end, we computed the delta-sample cell-cell interactions (ΔCCI) of the ligand-receptor pairs involving the shared PIGs for each aligned niche pair (Fig. 4i and Methods). *Apoe* encodes a canonical ligand of the receptor encoded by *Trem2* in the microglia, and this ligand-receptor pair facilitates Aβ uptake in the brain^26–28^. Our ΔCCI pattern of the ligand *Apoe* (Microglia) - *Trem2* (Microglia) receptor showed increased pattern and high spatial correlation with Aβ plaques in both 8-month and 13-month comparisons, which further validates the previous study^26–28^. However, the niches located in different regions showed diverse spatial patterns of the ΔCCI. For instance, the niches located in the cortical layer 4 (L4) and hippocampal CA1 (CA1) regions showed decreased interaction in the 8-month comparison, while the 13-month comparison niches exhibited increased interactions (Fig. 4i,j and Extended Data Fig. 6i), which implies that the crosstalk between *Apoe* and *Trem2* in these regions may be suppressed at early disease stages and be up-regulated in the later stages. When interrogating the ΔCCI patterns in the ligand *Apoe* (Astrocytes^29^) - *Trem2* (Microglia) receptor, we found that most of the hippocampus regions showed decreased interaction in the 8-month comparison and exhibited increased patterns in the 13-month comparison (Fig. 4i). This indicates that different cell type combinations present with the same ligand-receptor genes may show different ΔCCI patterns. Furthermore, we found that the ligand *Mfge8* (Astrocyte) - *Itgb5* (Microglia) receptor^30^ did not show a strong correlation to the Aβ plaque (Fig. 4i). By adding spatial and temporal patterns in ΔCCI analysis, our results may help further delineate the complex history of cell-cell interactions during disease progression.

Overall, the spatial gradient obtained through our ΔAnalysis reveals the heterogeneity of cell type abundance, gene expression, or cell-cell communications in diseased samples versus controls, which enables us to analyze disease pathology at a higher resolution.

### CAST Projection reconstructs spatially resolved single-cell multi-omics from multiple samples

Beyond performing ΔAnalyses, consistent spatial coordinates generated by CAST Stack further allow us to integrate samples with different spatial omic modalities. Here, we introduce CAST Projection, an unsupervised, label-free method to project single cells from query samples onto a reference sample toward spatially-resolved single-cell multiomics (Fig. 5a). To achieve this, it assigns single-cells from the query samples to the reference sample with the closest physical location and the most similar gene expression profile (e.g. the same cell type and cell state). In specific, we first conduct Combat^31^ and Harmony^32^ (Methods) single-cell data integration of the query and reference samples across different omic modalities to generate a shared low dimensional latent space, where cosine distance, a widely used metric in single-cell analysis^33–35^, is used to measure the similarity of cells across modalities. Given one reference cell, CAST Projection then searches for the cell with the closest cosine distance from the query sample within a confined physical radius as the matched cell pair (Methods). With well-aligned samples from CAST Stack, we can easily project the cells from multiple query samples to a shared reference sample with identical tissue coordinates.

**Fig. 5:**
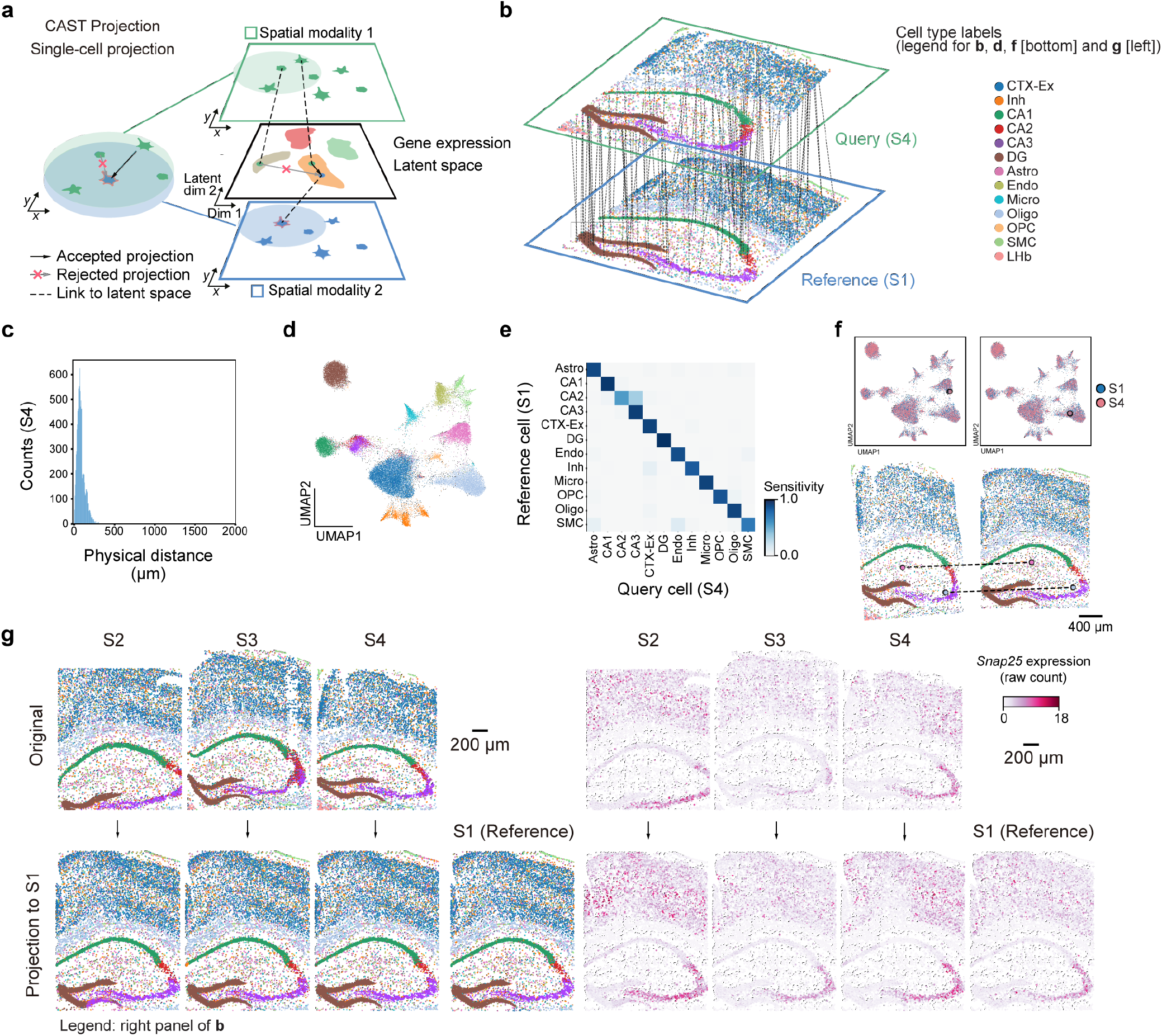
CAST Projection enables single-cell integration of spatial omics data across multiple samples. **a**, Strategy of the spatially and single-cell resolved cell assignment used by CAST Projection. Given physically aligned query and reference samples, CAST Projection first identifies candidate query cells for each reference cell within a radius based on cell density within the shared physical coordinate system; Next, it selects the query cell with the smallest cosine distance to the corresponding reference cell in the integrated gene expression latent space and assigns it to the reference cell. **b**, Schematic for CAST Projection results. Dashed lines (100 randomly sampled assignment pairs for visualization) connect cells from the query sample (S4) with its destination cell in the reference sample (S1). Colors represent cell types. **c**, The distribution of the physical distance in the spatial single-cell projection of the S4 to S1. **d**, Uniform manifold approximation and projection (UMAP) of the batch-corrected latent space across S1–S4 samples (control samples). Colors follow figure legends in **b. e**, Confusion matrix of the projection (S4 to S1, True Positive rate = 0.91). In order to analyze cell types with an adequate sample size, we filter out those that have less than 10 cells in the reference sample. **f**, CAST Projection assignment examples from S4 (query sample, pink) to S1 (reference sample, blue) in the UMAP plot (top) and spatial coordinates (bottom; colors follow figure legends in **b**). **g**, CAST Projection reconstructs one sample with multiple datasets. Left panel, the cells colored by cell types (colors follow figure legends in **b**); Right panel, the *Snap25* gene expression (raw count) profile in the original samples (top, S2–S4) and projected samples (bottom).

We first benchmarked the performance of CAST Projection using the four control samples (S1–S4) in the STARmap PLUS dataset^5^. When performing projection from S4 (query) to S1 (reference) (Fig. 5b and Supplementary Video 2), we evaluated the performance of CAST Projection by the Euclidean distances across all assigned cell pairs. The Euclidean distance of assigned cell pairs indicates that most of the cells in the query slice were assigned to the reference slice with small distances (Median distance = 72 μm; Fig. 5c). Meanwhile, we evaluated the accuracy of cell assignment using the cell-type labels (Fig. 5d and Extended Data Fig. 7a) between query-reference pairs — whether the query cell would be correctly assigned to the reference cell of the same cell type. As shown by the confusion matrix of cell type assignments (91% matched labels; Fig. 5e), the cell types of reference cells are highly concordant with their assigned query cells (Fig. 5f), which further supports that CAST correctly projects single cells from one tissue slice to another with the accurate match of spatial location and gene expression profiles.

Using CAST Projection, we finally integrated four biological samples (S1–S4) into one consistent spatial coordinate framework (Extended Data Fig. 7b-d) in which every single cell consists of 4 gene expression profiles from matched cells across samples S1–S4 (Fig. 5g). We further validated the accuracy of CAST Projection by checking the spatial distribution of gene expression profiles before and after projection, such as *Snap25* (Fig. 5g), *Mobp*, and *Tshz2* (Extended Data Fig. 7e), each of which showed consistent spatial patterns across S1–S4. As a result, CAST Projection successfully merged multiple slices into one common coordinate framework at the single-cell resolution^19^. Notably, experimental flaws (e.g. tissue distortion, slice fracture, missing imaging tiles) in individual slices do not significantly harm the performance of CAST and can be well compensated for by aggregating information from multiple samples through the spatial and single-cell integration of CAST Projection process (Fig. 5g). Furthermore, CAST Projection can also be applied across different major spatial omics technologies including Visium, MERFISH, Slide-seq, and Stereo-seq (Extended Data Fig. 7f and Supplementary Table 4).

### Spatially resolved single-cell translation efficiency analysis

To demonstrate the capability of CAST Projection to integrate different modalities of spatial omic measurements, we applied CAST Projection for four brain samples whose transcriptomes and translatomes were profiled respectively with STARmap and RIBOmap technologies at single cell resolution^7^ (Fig. 6a). While STARmap measures the cellular RNA expression with spatial information, RIBOmap selectively profiles the ribosome-bound RNA to probe protein translation *in situ*. The four-sample dataset is composed of two adjacent half brain slices taken from two different mice. Of the two adjacent slices, one was measured by RIBOmap and the other was measured by STARmap.

**Fig. 6:**
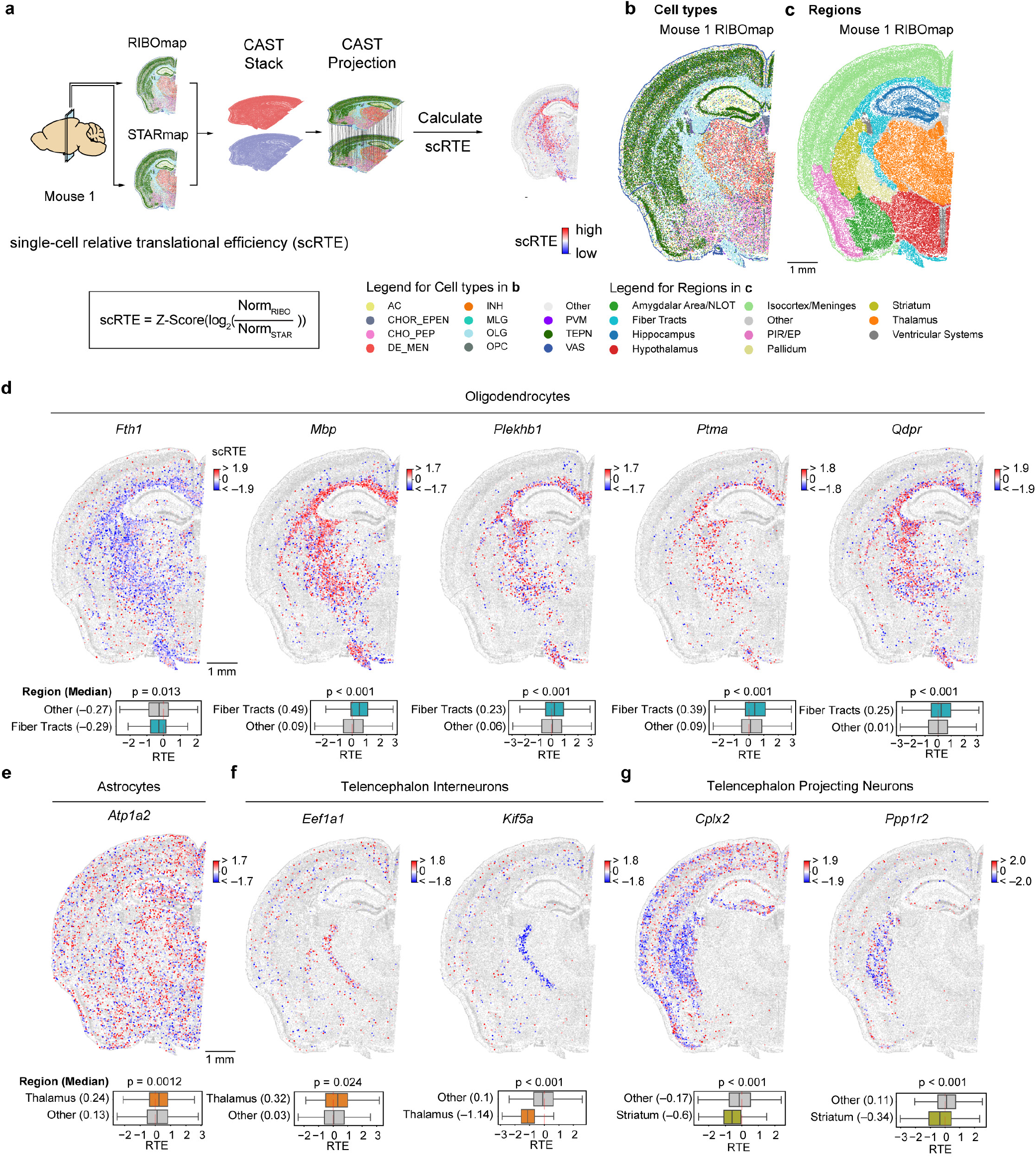
Single-cell resolved spatial-spatial integration of transcriptomics and translatomics reveals the ubiquitous heterogeneity of translation efficiency across cell types and brain regions. **a**, Schematic workflow for calculating single-cell relative translation efficiency (scRTE) profiles of the mouse brain. CAST Mark, Stack and Projection are sequentially performed on tissue samples to generate a spatially integrated sample where single cells have both RIBOmap and STARmap expression values. scRTEs are calculated using the formula. **b, c**, cell type and molecular tissue region profiles of the mouse1 RIBOmap sample (reference sample used in CAST Projection). AC, Astrocyte; CHOR_EPEN, Astro-ependymal cells; CHO_PEP, Cholinergic, monoaminergic and peptidergic neurons; DE_MEN, Di/Mesencephalon neurons; INH, Telencephalon interneurons; MLG, Microglia; OLG, Oligodendrocyte; OPC, Oligodendrocytes precursor cell; PVM, Perivascular macrophages; TEPN, Telencephalon projecting neurons; VAS, Vascular cells. **d-g**, Spatially resolved and cell type specific scRTE profiles in mouse1. Cells of the annotated cell type with available scRTE values are colored by scRTE levels, other cells are colored gray. The boxplots with Kruskal-Wallis test are used to evaluate the differences across the groups.

First, we performed joint cell typing and region segmentation using CAST Mark for the four brain samples, which resulted in 11 cell types (Fig. 6b, Extended Data Fig. 8a-c and Methods), and 10 major molecular regions as well as 23 sub-regions (Fig. 6c, Extended Data Fig. 8d,e and Methods). Next, we applied CAST Projection to project the STARmap cells to the RIBOmap cells after CAST Stack alignment (Fig. 6a). To validate the integration performance, we compared cell-type correspondence between query and reference cells, all of which showed accurate integration results (average percentage of matched labels = 85%; Extended Data Fig. 8f). After CAST Projection generated integrated tissue samples in which each cell contains both RIBOmap and STARmap measurements, we further defined single-cell relative translation efficiency (scRTE) as the normalized ratio of RIBOmap reads divided by STARmap reads in each cell (Fig. 6a and Methods). Although scRTE is not the absolute ratio of ribosome-bound RNA versus the total RNA as RIBOmap and STARmap were measured from two different samples using different technologies, it reflects the rank of relative translational levels compared to other cells in the dataset. We reason that scRTE is a more robust metric across samples while reflecting spatial heterogeneity of translation efficiency.

By profiling scRTEs across all genes, we sought to analyze the spatial heterogeneity of scRTEs across cell types and tissue regions. To this end, we first group genes into gene modules based on their mean expression profile across different cell types, which results in 11 gene modules (M1-M11, Extended Data Fig. 9a,b). We then conducted cell-type specific scRTE analysis within each cell type with gene modules that have adequate expression: M1–M5 and M9 in neurons, M6 in astrocytes, M7 in microglias, M8 in oligodendrocytes, M10 in vascular cells and M11 in Astro-ependymal cells (Supplementary Table 7), which revealed widespread cell-type- and tissue-region-dependent translational regulation.

In oligodendrocytes, we detected dramatically different scRTE levels of M8 genes between fiber tracts and other regions, which involve axon ensheathment, nervous system development, and myelination. For example, *Mbp, Plekhb1, Ptma* and *Qdpr* show significantly higher scRTE levels in fiber tracts, in contrast, *Fth1* shows relatively lower scRTE levels (Fig. 6d and Extended Data Fig. 10a). The differential translational regulation of these genes in the fiber tracts versus other regions indicates regional specialization of protein synthesis to support oligodendrocyte functions (e.g. myelination). In astrocytes, the *Atp1a2* shows higher scRTE levels in the thalamus region (Fig. 6e and Extended Data Fig. 10b). In telencephalon interneurons, the translation elongation factor *Eef1a1* has higher scRTE levels in thalamus than other regions, while the *Kif5a* exhibits lower levels in thalamus (Fig. 6f and Extended Data Fig. 10c). In the telencephalon projecting neurons, *Cplx2* and *Ppp1r2* both show lower levels in the striatum region (Fig. 6g and Extended Data Fig. 10d). These results support the heterogeneity of translation efficiency across different cell types or anatomical regions, and the necessity to investigate mRNA translation regulation with both single-cell and spatial resolutions in future studies.

## Discussion

In summary, we demonstrated that CAST enables search-and-match across samples based on their spatially resolved molecular similarities while uncovering and visualizing the variability driven by spatial differences. CAST is capable of integrating the samples across different technologies and different spatial modalities at single-cell resolution. All three CAST modules only require gene expression and spatial co-ordinates of cell/voxels, without the need of cell type labels, histology, or DAPI images that are not available for all datasets, suggesting a wider applicability than existing methods to current and future datasets (Supplementary Table 1,4 and 5). CAST has broad applicability in major spatial omics technologies including Visium, MERFISH^36^, Slide-seq^37^, Stereo-seq^4^, STARmap and RIBOmap (Supplementary Table 4,5). Notably, in a single run, CAST successfully aligned three half brain samples collected from MERFISH, Slide-seq, and STARmap, respectively (Fig. 3a). The multi-technology (3 and above) alignment enabled by our method will be an essential integration tool for large-scale tissue datasets across multiple labs towards molecularly defined common coordinate framework (CCF)^19^ of biological tissues. In addition, CAST is also applicable to variable gene panel sizes (from 64 to genome-wide) with as few as 64 overlapping genes, which shows great promise to further integrate the spatial omics data with limited feature panel sizes across a wider scope of spatial technologies, such as antibody-based spatial proteomics data. Such multi-technology spatial-spatial integration will benefit users to combine the strengths of different spatial technologies by cross-reference across various spatial resolution and gene panels.

To demonstrate the downstream application of aligning different spatial omic samples into consistent tissue coordinates, we introduced ΔAnalysis, a computational framework for spatial differential analyses across disease conditions, in compliance with the widely used Anndata data structure^38^. Using ΔAnalysis, we discovered the spatial heterogeneity of the different molecular characteristics in AD, such as gene expression, cell type abundance, and cell-cell communication, which opens new perspectives toward a deeper understanding of disease mechanisms. We also integrated spatially resolved *in situ* ribosome-bound RNA profiling (RIBOmap) and RNA profiling (STARmap) to uncover the spatial translation efficiency landscape of brain tissues at the single-cell level using the scRTE metric. We found that populations of single cells of one cell type may show heterogeneous gene-specific translation efficiency across different regions.

In summary, CAST provides a comprehensive and modular framework for the integration and differential analyses of spatial omics data across biological replicates, measurement modalities, and disease conditions with both spatial and single-cell resolutions.

## Methods

### Data preprocessing

In all the spatial omics datasets used, we normalized the sum of the raw read counts of each cell to1×10^4^ (referred to as *norm1e4*). We then applied a log2-transformation to the normalized counts (referred to as *log2_norm1e4*). Finally, the expression values were scaled without zero-centering (referred to as *scale*). Each data transformation is stored as an Anndata layer.

### CAST Mark algorithm

Given a sample with cells, the corresponding dataset is composed of each cell’s spatial co-ordinates Ψ ∈ ℝ^*M*×2^(x and y coordinates) and the feature expression matrix *X* ∈ ℝ^*M*×*N*^ (*N* indicates the feature dimension, e.g. gene expression panel size). For each tissue sample, we first constructed the tissue graph by performing Delaunay triangulation using the spatial coordinates, resulting in an adjacency matrix *A* ∈ ℝ^*M*×*M*^.

The CAST Mark GNN is composed of with *L* GCNII layers^15^ after an optional single-layer perceptron encoder. The perceptron encoder serves as an option to reduce feature dimension and thus reduces the demand for computational resources without large compromise in performance. For each layer *l*(*l* = 1, 2, …, *L* − 1),

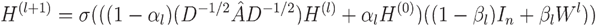

Where σ(·) is a non-linear activation function (by default, ReLU). *Â* is the adjacency matrix with self-loops, *D* is its diagonal degree matrix. H^(0)^ is the initial node features (e.g. gene expression for each cell), while H^(*l*)^ is the feature for layer *l. α*_*l*_ and *β*_*l*_ are hyperparameters for which we used their default values in the DGL package^39^.

We utilized a self-supervised CCA learning objective^40^ to train the network, where for each sample, we first applied random node feature masks and random edge masks to the initial graph *G* to generate two augmented views of 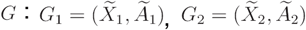 providing a mechanism to tolerate the intrinsic or sample-level stochasticity of gene expression and spatial locations of cells at microscopic scales. The CAST Mark GNN *∈*_*Θ*_ is subsequently employed in parallel to create node embeddings for the two augmented views: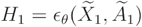 and 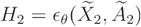 Then we normalized 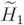 and 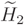 by

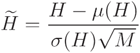

Where *μ* is the mean value of each feature in the given matrix, σ indicates the standard deviation of the values in each feature, and *M* is the number of the cells. The normalized 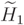 and 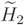 are used for the CCA-based self learning objective. The objective function is:

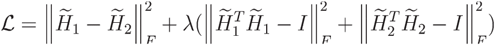

where the *I* is the identity matrix.

In this study, we used *L* = 9 for all analyses. After the training process, the final graph embedding of the original graph *G* is 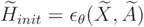.

#### Performance evaluation

We used Pearson correlation to evaluate the similarity of the graph embeddings. We used the adjusted rand index (ARI) and the percentage of consistent cells between corresponding clusters to evaluate the clustering performance.

### CAST Stack algorithm

To align the spatial coordinates of samples while preserving the cell organization, CAST Stack performs alignment using a gradient-descent-based rigid alignment phase followed by a non-rigid alignment phase to achieve a proper transformation.

#### Rigid Alignment

Affine transformation is used for rigid registration. CAST provides translation, rotation, scaling and reflection transformations, but disallows shear mappings. With the coordinates Ψ = (*x*_*i*_, *y*_*i*_), *i* = 1, …, *M* of the *M* cells in the query sample, the affine transformation algorithm can be written as:

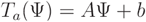

Where *T*_*a*_ is the affine transformation function with the transformation matrix *A* and the translation vector *b*:

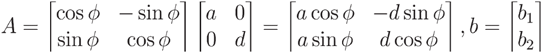

To automatically find a proper transformation, a gradient descent (GD) optimizer was performed for the affine transformation. We identify the *θ*as vector of the five parameters *a,d, ϕ,b*_1_ and *b*_2._:

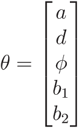

The cost function *J* is identified as the sum of the Pearson distance *J*_i_ between each query cell *i* and its nearest reference cell:

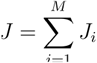

Optimization steps are formulated as:

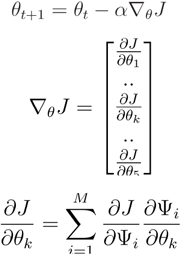

where *α* is a weighting parameter of the gradient descent.

The 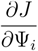 is the partial derivative of the with respect to coordinate variable Ψ_i_:

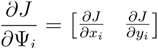

The 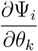 is the partial derivative of the coordinate variable Ψ_i_:with respect to*θ*_*k*_ :

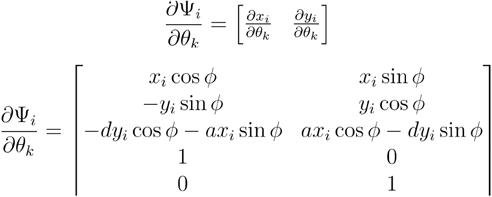

#### Non-rigid Alignment

The free-form deformation (FFD) based on the B-splines method is used for the deformable transformation^41^. To define a spline-based FFD, we first generated the mesh grid for the spatial slice. Given the number of the control points *L* in each dimension, the mesh spacings *g*_*x*_ and *g*_*y*_ are calculated by:

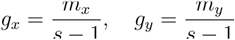

where *m*_*x*_ and *m*_*y*_ represent the maximum coordinate of the slice. Ω indicates a *s* × *s* control points ω_*i,j*_ in the mesh grid with spacing *g*_*x*_, *g*_*y*_, respectively. All the cells (*M* cells) in a given query sample are regarded as Ψ = {(*x, y*)|0 ≤ *x* ≤ *m*_*x*_, 0 ≤ *y* ≤. *m*_*y*_}. The B-Spline transformation matrix *T*_*b*_ for each control point is written as:

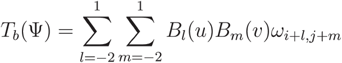

where *i* = ⌊ *x*/*g*_*x*_⌋, *j* = ⌊,*y*/*g*_*y*_⌋, *u* = *x* − *i* ∗ *g*_*x*_, *v* = *y* − *j* ∗ *g*_*y*_, and where *B*_*l*_, *B*_*m*_ represents the *l*-th and *m*-th basis function of the B-spline, respectively:

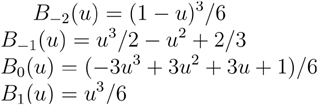

Similarly, the formula of the gradient-descent-based FFD is written as:

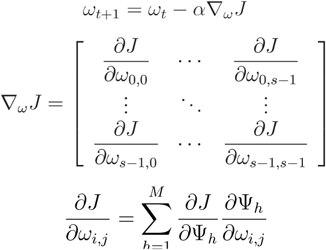

where *α* is a weighting parameter of the gradient descent.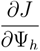 is the partial derivative of the with respect to coordi-nate variable Ψ_*h*_ :

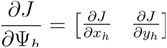

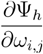 is the partial derivative of the coordinate variable Ψ_*h*_ with respect to ω_i,j_, which is equal to 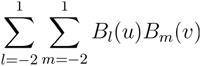.

For efficient computation of the samples with a large number of cells (such as the half-brain samples), we also provide an optional downsample strategy. CAST Stack could select a random subset of cells (e.g. 20,000 cells) and perform CAST Mark or Stack on the dataset at a reduced size. After generating the transformation parameters, we then apply the calculated alignment transformation to all cells in the original sample at the full resolution.

### CAST Projection algorithm

We assume that a given cell will be most similar to the cells with close distance in physical space and low-dimensional feature space. Thus, to project the features of the cells into a low-dimensional space, CAST Projection employs a sequential combination of Combat^31^ and Harmony^32^ integration for samples with different modalities. Cosine distance is used to measure the similarity of cell features in the integrated embedding. To find the candidate cells for a given reference cell, CAST first identifies the candidate query cells within a radius of the reference cell. As different cell types exhibit varying cell distances in the space, CAST calculates the cell-type specific cell average distance based on the Delaunay triangulation graph. By default, twice the averaged distance is utilized (In AD samples, 1.5 times cell distance is used, while in RIBOmap-STARmap, three times the distance is used). Among the candidate query cells, CAST identifies the *k* cells with the closest cosine distance to project. Inverse distance weighting (IDW) algorithm is used to interpolate the weighted average of the features in *k* candidate query cells as the feature of the projected cell. The interpolated gene expression 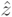is

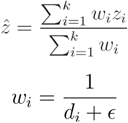

Where the *ω*_*i*_ is the interpolation weight, *d*_*i*_ is the cosine distance between the reference cell and *i*-th candidate query cell, *∈* is a hyperparameter to adjust the extreme weight.

In this study, we set *k* = 1, which means that the projection task is equivalent to the cell assignment.

### Simulation Datasets

To generate a dataset with ground truth cell partners across samples, we took S1 from the STARmap-PLUS AD dataset as our reference, and generated one simulated sample based on S1 where each cell in the synthetic sample corresponded to a ground truth partner in the S1 sample. The simulated sample was generated by the following steps:

1. Physical location noise (non-linear): Gaussian Process Warp^42^ was used to perturb the spatial coordinates of the reference sample using the following parameters: slope_variance = 1e-3; noise_variance = 1e5; kernel_variance = 1e5; kernel_lengthscale = 1.0; mean_slope = 1.0; mean_intercept = 0.1.
2. Global spatial coordinates distortion (linear): The tissue sample was further changed by scaling and rotation transformations (x-axis: 40%; y-axis: 50%; rotation: 30º)
3. Gene expression noise: We applied gaussian noise (*μ* = 0, *σ* = 0.2) to the *log2_norm1e4* gene expression matrix.
4. Gene feature dropout: We randomly replaced 10% of the values in the expression matrix using zeros.
5. Cell dropout: We randomly dropped 10% of cells in the simulated sample, making sure that graph structures will be altered.

### Region marker gene detection

We calculated the average gene expression (*log2_norm1e4*) in each region, which represents the gene expression abundance. Then, Z-scores of these averaged values are calculated across all regions to quantify the degree to which expression levels vary across different regions^43^. By considering these two features and comparing them with the databases^17^, we identify the region marker genes (Supplementary Table 3) with expert curation.

### Delta-sample analysis

The delta-sample analysis (ΔAnalysis) is used to discover the variance driven by spatial differences across conditions. With the well-aligned samples, given one neighborhood (niche), we could get the cells and their molecular characteristics in this neighborhood with different conditions. For each cell, we define a neighborhood as all the neighboring cells within a default 50 μm radius of its center. By comparing the associated neighborhoods of aligned cells, we obtain delta statistics for molecular features such as gene expression and cell type abundance at a local resolution on the global tissue slice. After screening all cells in the sample, we obtain a global spatial gradient map of the differences in molecular features between conditions. In this study, we use these molecular features in each neighborhood:

#### Cell type abundance

the cell count of a certain cell type. The ΔCell is the difference of the cell type abundance in each comparison. For example, for the one of the combinations (S8 - S3) in the 13-mos comparison.

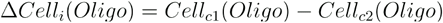

where is the *cell*_*c*1_(*Oligo*)abundance of the oligodendrocytes in disease samples S8, while *cell*_*c*2_(*Oligo*) is the abundance of the oligodendrocytes in control samples S3. The strategy is applied for *Gene expression, Cell-cell adjacencies* and *Cell-cell interactions*.

#### Gene expression

The ΔExp is the difference of the average gene expression (*log2_norm1e4)* in one cell type in each comparison. The spatial PIGs are identified by the following criteria: (1) ΔExp > 0.1; (2) The spatial correlation (Pearson’s Correlation) between the ΔExp and plaque score is greater than 0.1; (3) The FDR values of the Wilcoxon rank sum test for the differential expression analysis is less than 0.1.

#### Cell-cell adjacencies

The ΔCCA is defined as the difference of the cell-cell adjacency value of the given cell type pairs. The cell-cell adjacency value between cell type A and B is defined as the number of A-B edges within a neighborhood on the tissue graph.

#### Cell-cell interactions

The ΔCCI is the difference of the cell-cell interaction degree of a ligand-receptor pair in each comparison derived as in CellPhoneDB^44^. The cell-cell interaction degree is calculated by Squidpy (1.2.2)^45^ with the normalized counts (*norm1e4*).

#### Plaque score

the sum of the plaque area in a given niche. We filtered plaques with the area less than 300 pixels (∼ 30 μm^2^) in the image.

#### Tau score

the sum of the tau rate in the cells. The tau rate is defined as the ratio between the tau area and the cell area in each cell.

To interrogate the spatially resolved molecular differences among different age groups, we use two comparisons: 8-month disease and control (8 mos), 13-month disease and control (13 mos; Fig. 4a).

### Single-cell Relative translation efficiency (scRTE) analysis

In order to measure the translation efficiency among cells regardless of the different expression distributions due to the different technologies or samples, we introduce the scRTE metric for each cell as the following formula:

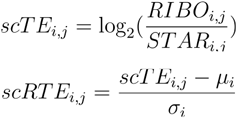

Where *RIBO*_*i,j*_ and *STAR*_*i,j*_ are the RIBOmap and STARmap normalized counts (*norm1e4*) of the gene *i* in cell *j*. The *μ*_*i*_ and *σ*_*i*_ are the average value and standard deviation of the *scTE*_*i,j*_ of the gene *i* across all cells. The *scRTE*_*i,j*_ is the z-score of the *scTE*_*i,j*_ over all cells.

Once we calculate the scRTE values of each cell in a given gene, scRTE levels at different locations may not be consistent. To detect the spatial variability of the scRTE levels in each gene, we use the standard deviation of the scRTE values of each gene to measure the degree of heterogeneity for each gene. Meanwhile, the Kruskal-Wallis test is used to evaluate whether the scRTE levels are significantly different between the cell types or regions. As the STARmap sample in Mouse 2 is truncated at the hypothalamus, cortical subplate and olfactory cortical regions, our analysis focuses solely on the overlapping region within the Mouse 2 sample.

### Sample preparation of STARmap sample (mouse 2)

The mouse (C57BL/6 strain) utilized in this study was anesthetized with isoflurane and subsequently decapitated quickly. We collected the brain tissue and embedded it in the Tissue-Tek O.C.T. Compound, which is next frozen in liquid nitrogen and stored at -80 °C. The mouse brain tissue was further transferred to a cryostat (Leica CM1950) at -20°C and was sliced into 20 μm coronal sections. These slices were then placed on glass-bottom 12-well plates that had been pre-treated with 3-(Trimethoxysilyl)propyl methacrylate and poly-D-lysine. Next, the slices were fixed with 4% PFA in PBS for 15 minutes (room temperature), permeabilized with cold methanol and kept at -80 °C for an hour. The experimental procedures for STARmap were similar to those used for HeLa cells in Zeng et al.^7^, except that all the reaction volumes were doubled because the brain tissue was prepared in 12-well plates. We captured the images using a Leica TCS SP8 confocal microscopy with a 63X oil immersion objective (NA 1.4) and a voxel size of 90.14 nm X 90.14 nm X 300 nm. In the first sequencing round, we also imaged the DAPI staining signals. Overall, 9 imaging cycles were sequenced to detect the 5,413 genes.

### Data processing for STARmap brain tissue sample

#### STARmap Imaging Preprocessing

We used Huygens Essential version 21.04 (Scientific Volume Imaging, The Nether-lands, http://svi.nl) to deconvolute the raw images (CMLE algorithm ; SNR:10 and 10 iterations). The image preprocessing operations, such as image registration, spot calling, and barcode filtering with minor adjustments were applied following the previous reporting^7^.

#### Cell Segmentation

The RNA amplicon based cell segmentation method ClusterMap^24^ was utilized to automatically detect the cells. We applied the default pipeline with minor adjustments to the DAPI signal preprocessing in order to capture cells in each field of view (FOV). The following parameters were used in the ClusterMap: cell_num_threshold = 0.7; dapi_grid_interval = 5; pct_filter = 0.1; window_size = 550. After the cell segmentation for each FOVs, cells were stitched to generate the cell-by-gene matrix.

#### Quality control and Cell typing

We adopted the quality control strategy reported by Zeng et al.^7^ the STARmap (mouse 2) sample. The median absolute deviation (MAD) was used to estimate the lower/upper boundaries for cell filtration:

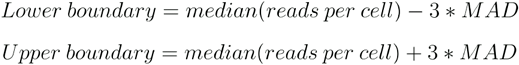

The cells with the reads fewer than lower boundary or greater than upper boundary were filtered out, resulting in 44,751 cells and 5,413 genes. Employing the similar strategy reported by Zeng et al.^7^, we further identified 11 cell types across 4 samples.

### Region Segmentation of mouse half brain datasets

We first performed CAST Mark training on the normalized expression (*norm1e4*) with Combat batch correction^31^ of 1,082 highly variable genes across all 4 half-brain samples. We then performed *k*-means (k = 20) clustering on the CAST Mark graph embedding. Among the 20 clusters, we selected the most under-segmented cluster (region 3) and further sub-clustered region 3 into 10 sub-clusters, yielding a total of 29 clusters. We then visually examined all 29 regions. Using the Allen Brain Atlas^17,18^ as the reference, we merged over-segmented regions consistent with established brain anatomy. We also separated physically segregated areas belonging to the same *k*-means cluster (HY and LH). Consequently, we confirmed a total of 23 brain subregions. Finally, we concluded these 23 brain subregions into 10 top-level brain regions based on the Allen Brain Atlas.

### Gene Clustering and Enrichment Analysis

The gene expression (*log2_norm1e4*) of the 4 samples were first averaged across the cell types within each sample, respectively. Subsequently, the average expression values were standardized by calculating the Z-score within each sample. The 884 highly abundant genes with sufficient expression and scRTE values in each sample were used in this analysis. The standardized vectors were merged and clustered with the Louvain algorithms from Seurat (Version 4.0.3). We then used ComplexHeatmap (Version 2.10.0) to visualize the clusters. To identify the enriched GO and KEGG pathway terms, gprofiler2 (Version 0.2.1) was applied to do the enrichment analysis. The enriched terms were further visualized by the EnrichmentMap plugin in Cytoscape (Version 3.9.1). For visualization, clusters containing fewer than five nodes are excluded. For the spatially-resolved PIGs, the GO and KEGG pathway enrichment analyses are conducted with clusterProfiler (Version 3.18.1)^46^.

### Benchmark with PASTE Alignment

We use the pairwise_align (GPU) and center_align (CPU mode; not available in GPU mode) in PASTE to run the alignment tasks of different samples with default parameters. The NVIDIA RTX A5000 (24GB VRAM) is used in the task. We only present the results for the 8 AD samples, as PASTE was unable to execute the half-brain alignment tasks (limited to CPU 80 GB RAM).

## Supporting information

STable1

STable2

STable3

STable4

STable5

STable6

STable7

## Data availability

The RIBOmap and STARmap datasets are available from (RIBO-map_mouse1, STARmap_mouse1 and RIBOmap_mouse2: https://singlecell.broadinstitute.org/single_cell/study/SCP1835; STARmap_mouse2: https://singlecell.broadinstitute.org/single_cell/reviewer_access/d7977efa-b46a-4acf-9372-a5fec45bdea9). The AD STARmap PLUS dataset is publicly available at (https://singlecell.broadinstitute.org/single_cell/study/SCP1375/).

Two Visium datasets (Mouse Brain Coronal Section 1 (FFPE) and Mouse Brain Coronal Section 2 (FFPE)) used are available from:

https://www.10xgenomics.com/resources/datasets/mouse-brain-coronal-section-1-ffpe-2-standard

https://www.10xgenomics.com/resources/datasets/mouse-brain-coronal-section-2-ffpe-2-standard

The MERFISH dataset used (co1_slice37 in co1_sample13) is available from:

https://doi.brainimagelibrary.org/doi/10.35077/act-bag

The Slide-seq dataset used (slice042) is available from:

https://docs.braincelldata.org/downloads/index.html

The two Stereo-seq MOSTA datasets used (E16.5_E2S5 and E16.5_E2S6) are available from:

https://db.cngb.org/stomics/mosta/download/

## Code availability

The code and demos of the CAST have been deposited to Github (https://github.com/wanglab-broad/CAST). The implementation of CAST, as well as the tutorials, is available in the demo pipeline files.

## Acknowledgements

We thank Hailing Shi and Yiming Zhou for their help in brain regions, Haowen Zhou, Kamal Maher, Jiakun Tian, Wenbo Wang, and Peng Tan for discussion. Z.T. thanks Xiao Jin for his guidance in formulating the algorithms, Yuan Zhou for technical assistance. S.L. thanks Wentao Mo for the discussion on graph neural networks. X.W. gratefully acknowledges support from the Searle Scholars Program, Thomas D. and Virginia W. Cabot Professorship, Edward Scolnick Professorship, Ono Pharma Breakthrough Science Initiative Award, Merkin Institute Fellowship, and NIH DP2 New Innovator Award.

## Author contributions

X.W., Z.T. and S.L. conceived the study. Z.T. and S.L. formulated, developed, and implemented the CAST algorithm. H.Z. collected RIBOmap and STARmap half brain data. Z.T. and J.H. performed data preprocessing of the RIBOmap and STARmap half brain data. J.H. reproduced CAST code. Z.T. and S.L. performed data analysis, under the supervision of X.W. The manuscript was written by Z.T., S.L., M.W., and X.W. All authors read and approved the manuscript.

## Competing interest statement

X.W. is a scientific co-founder of Stellaromics. X.W. and H.Z are inventors on pending patent applications (International Application No. PCT/US2022/031275, PCT/US2022/035271) related to STARmap PLUS and RIBOmap. Other authors declare no competing interests.

**Extended Data Fig. 1:**
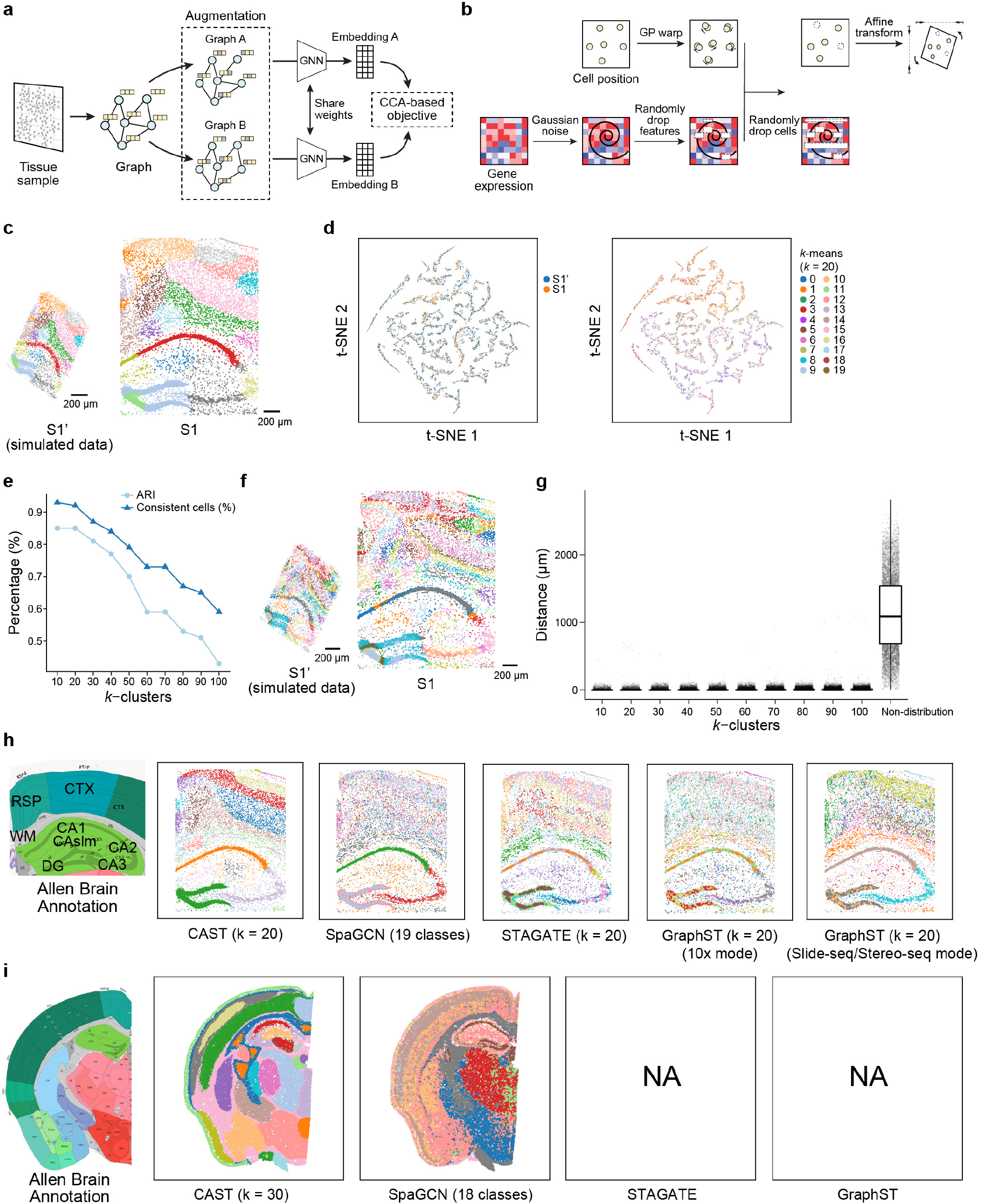
CAST Mark identifies the common spatial features between the simulated and real samples. **a**, The schematic workflow of the self-supervised learning framework used in CAST Mark. **b**, The simulation strategy to generate the simulated dataset S1’ from the real sample S1 (8 month, control) in STARmap PLUS dataset (Methods). The gene expression matrix of the S1 sample is perturbed by adding gaussian noise and randomly dropping features and cells. The Gaussian Process warping42 and the affine transformation are applied to distort the coordinates of S1. **c**, The *k*-means (*k* = 20) result of the graph embedding generated by CAST Mark. The different colors in the cells indicate the different clusters of the graph embedding. **d**, The t-distributed stochastic neighbor embedding (t-SNE) visualization of the spatial embedding labeled with samples (left) and *k*-Means clusters (*k*=20, right). **e**, The clustering performance (Adjusted Rand Index and the percentage of the consistent cells) in different numbers of the clusters (*k*). The percentage of the consistent cells indicates the percentage of the cells in two datasets (simulated sample S1’ and real sample S1) are classified into the same cluster across both samples. **f**, The *k*-means (*k* = 100) result of the spatial embedding generated by CAST Mark. **g**, The distance distribution of the cells in different *k*-clusters and non-distribution groups. The distance indicates each cell in the simulated sample S1’ to the closest one with the same cluster in the S1 sample. Average distance: 2.16 μm (*k* = 10), 3.25 μm (*k* = 20), 4.46 μm (*k* = 30), 5.21 μm (*k* = 40), 6.43 μm (*k* = 50), 8.02 μm (*k* = 60), 8.39 μm (*k* = 70), 9.51 μm (*k* = 80), 10.54 μm (*k* = 90), 11.53 μm (*k* = 100) and 1120.93 μm (Non-distribution). **h**, Enabled by a deep GNN and a self-supervised CCA objective, CAST Mark outperforms existing GNN-based methods (SpaGCN, STAGATE, GraphST) in tissue segmentation tasks at a ∼9,800-cell scale. Segmentation results were colored by clusters, shown along with brain annotation from the Allen Institute17,18. **i**, Tissue segmentation was performed on a STARmap dataset collected on a single coronal slice of mouse half brain7. The performance of CAST Mark scales up to the ∼60,000-cell scale (a typical coronal slice of mouse half brain), outperforming SpaGCN11, while GraphST and STAGATE fail to handle this large dataset at single-cell resolution.

**Extended Data Fig. 2:**
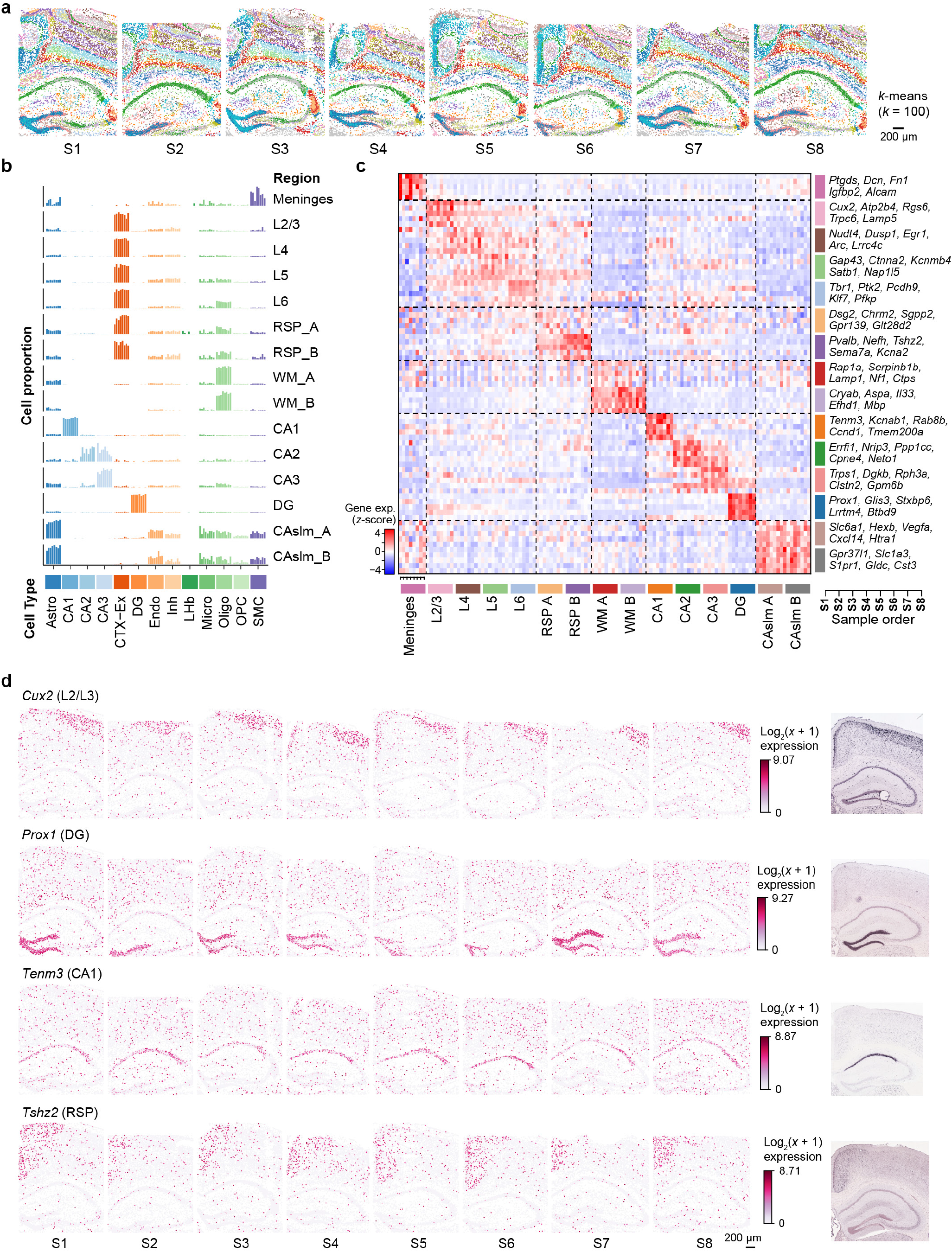
CAST Mark identifies consistent regions across age, strain, and disease conditions. **a**, The *k*-means (*k* = 100) clustering result of graph embeddings generated by CAST Mark in the samples S1–S8. Different colors in the cells indicate the different clusters of the embedding. **b**, The bar plots show the consistent cell proportions of each cell type in each sample (Ordered from S1 to S8) and region. Astro, Astrocyte; CA1, CA1 excitatory neuron; CA2, CA2 excitatory neuron; CA3, CA3 excitatory neuron; CTX-Ex, Cortex excitatory neuron; DG, Dentate Gyrus; Endo, Endothelial cell; Inh, Inhibitory neuron; Micro, Microglia; Oligo, Oligodendrocyte; OPC, Oligodendrocyte precursor cell; SMC, Smooth muscle cell. **c**, The Z-score of the mean value of the *log2_norm1e4* (Methods) profile of region marker genes (Supplementary Table 3) in each region (Ordered from S1 to S8) shows that CAST Mark identifies the consistent regions across multiple samples. **d**, The gene expression of region markers *Cux2* (L2/L3), *Prox1* (DG), *Tenm3* (CA1) and *Tshz2* (RSP) across 8 samples (Ordered from S1 to S8). The region marker genes are validated by the ISH images from the Allen Brain Atlas17.

**Extended Data Fig. 3:**
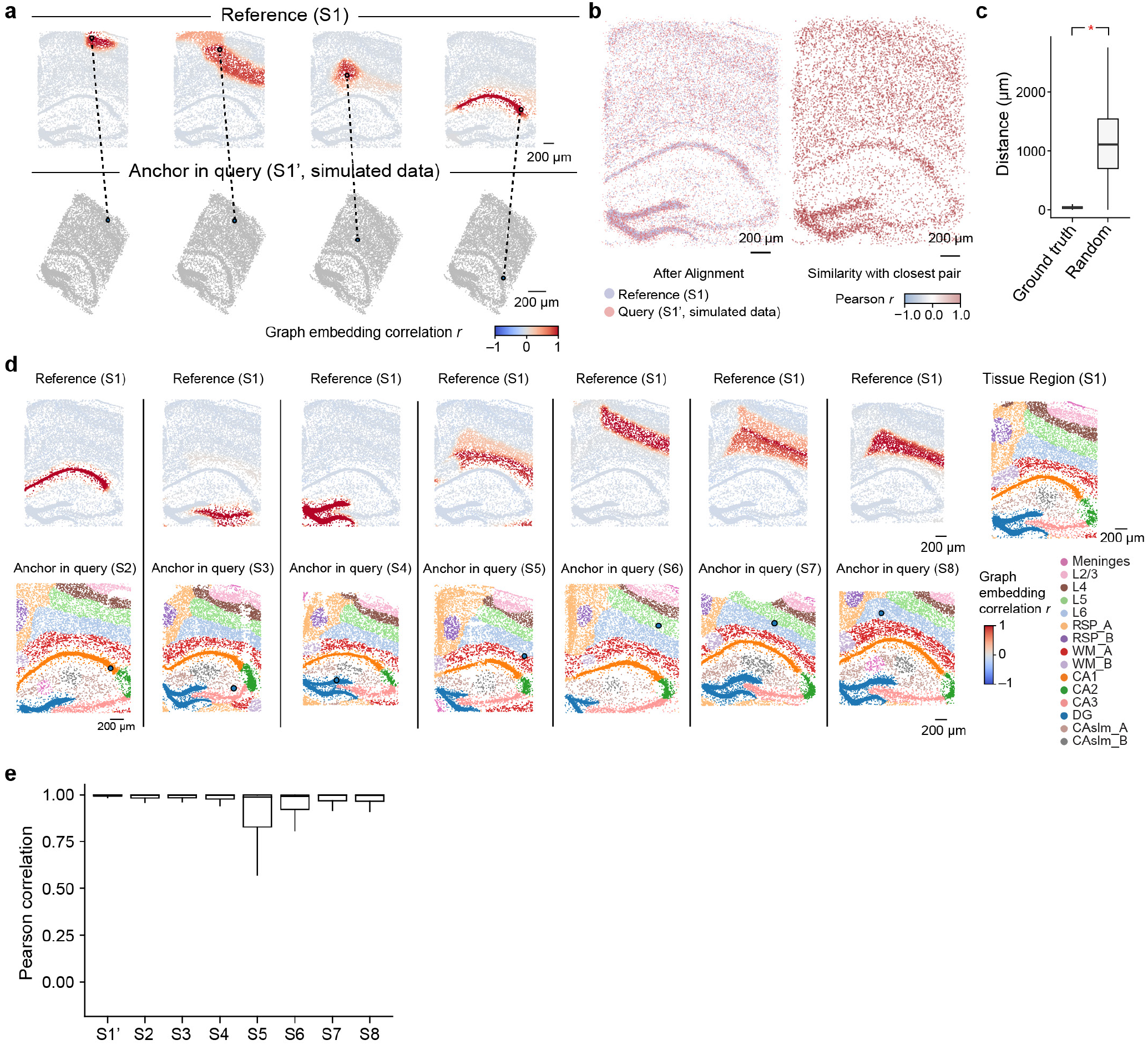
Pearson correlation of CAST Mark graph embedding between cells is a robust similarity metric for cell locations across samples. **a**, CAST Stack can predict relative cell locations in the synthetic dataset. Given the query cell in the query slice (simulated dataset S1’), the cells in the reference sample (S1) are colored by Pearson’s correlation of the graph embedding between the reference cell and the given query cell in the query sample (S1’). **b**, Left panel, the coordinates of the S1 and the S1’ after alignment. Right panel, each cell in the query sample is colored by Pearson correlation of the graph embedding between itself and its closest pair in the reference sample. **c**, The boxplots show the significant closer physical distances (One-way ANOVA) of the correct cell pair (ground-truth, mean = 37.82 μm, sample size = 8,789) than the ones in random cell pairs (Random, mean = 1133.49 μm, sample size = 8,789). **d**, CAST Stack can map cells to relative tissue regions in real tissue datasets. Given a query cell in the query sample (S2–S8), the cells in the reference sample (S1) are colored by Pearson’s correlation of the graph embedding between the reference cells and the given query cell. Cells with high Pearson’s correlation in the reference sample show similar relative spatial locations to the query cell. In the bottom panel, cells in the query samples are colored by tissue region labels produced by CAST Mark. **e**, The boxplots show the high Pearson correlation of the graph embedding between the cells in query samples (S1’ and S2–S8) and the reference cell with the closest physical distance in the reference sample (S1). Average Pearson *r*: 0.99 (S1’, *n* = 8,789), 0.95 (S2, *n* = 8,506), 0.94 (S3, *n* = 9,428), 0.93 (S4, *n* = 8,034), 0.82 (S5, *n* = 8,202), 0.88 (S6, *n* = 8,186), 0.91 (S7, *n* = 9,634) and 0.92 (S8, *n* = 10,372).

**Extended Data Fig. 4:**
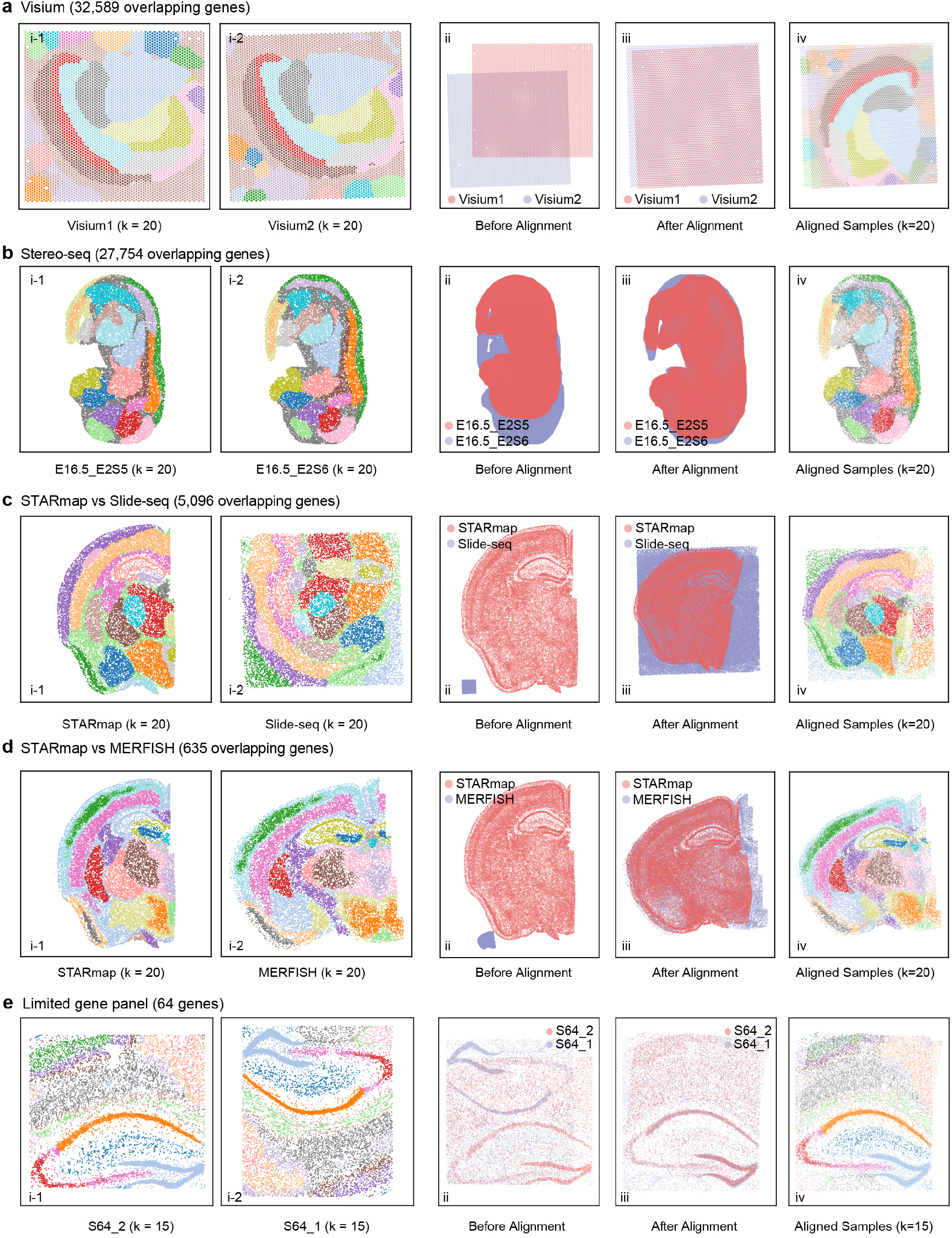
CAST has wide utility across various spatial technologies. **a-e**, CAST automatically searches and matches shared tissue anatomy in different technologies at high spatial resolution: **a**, Visium (Visium1: Mouse Brain Coronal Section 1; Visium2: Mouse Brain Coronal Section 2); **b**, Stereo-seq; **c**, STARmap (STARmap_mouse1) vs. Slide-seq; **d**, STARmap (STARmap_mouse1) vs. MERFISH; **e**, Two STARmap samples with limited gene panels (both have 64 genes). **i**, joint *k*-Means clustering results of CAST Mark graph embeddings of two samples, colored by joint clusters. **ii**-**iii**, spatial coordinates of the query sample (colored pink) and the reference sample (colored blue) before (**ii**) and after (**iii**) alignment. **iv**, aligned samples colored by joint *k*-Means clustering results of the graph embeddings.

**Extended Data Fig. 5:**
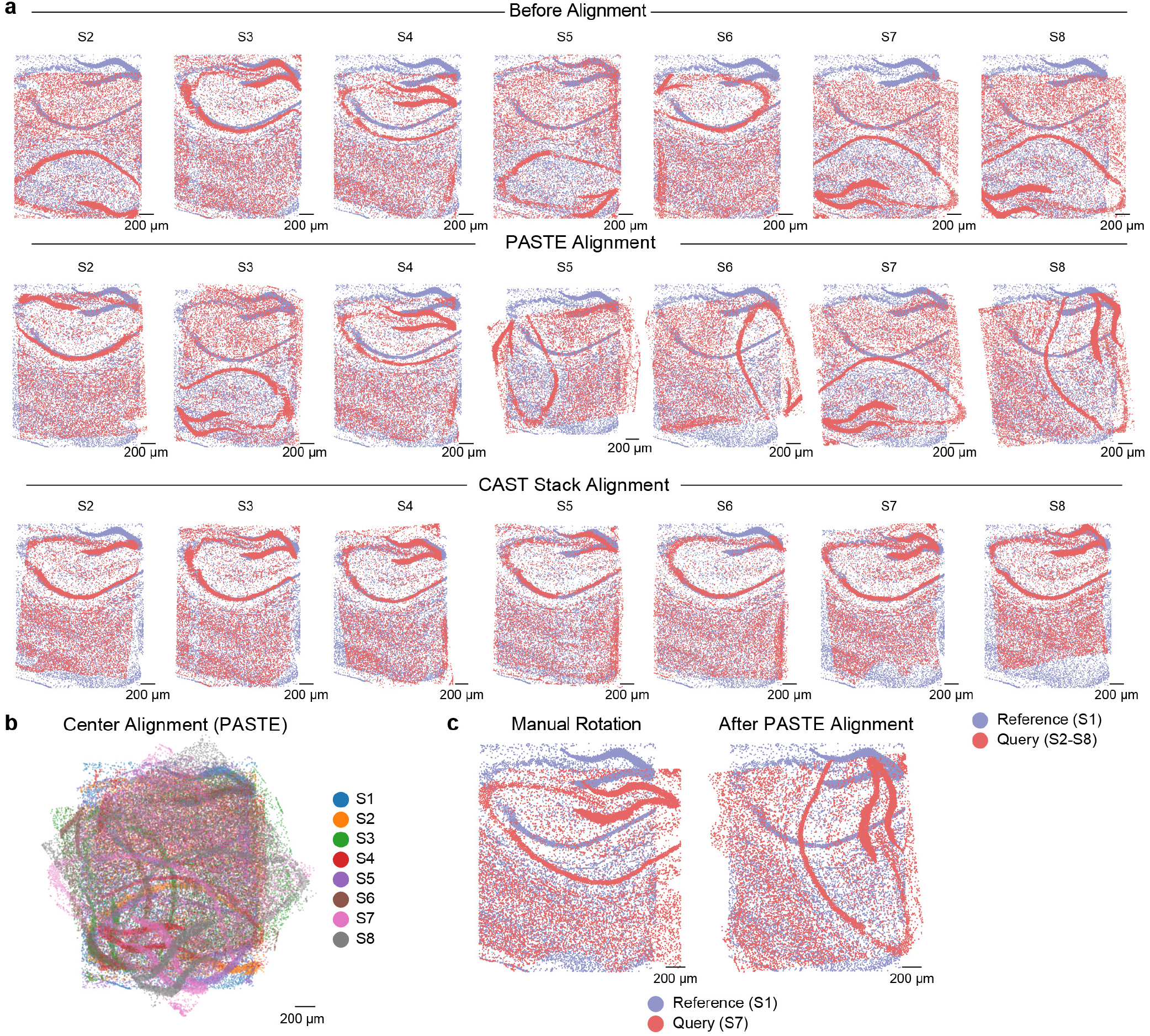
CAST Stack outperforms existing alignment methods. **a**, Benchmarking the performance of CAST and PASTE in one-to-one alignment tasks. Spatial coordinates of query (S2-S8, respectively, colored in pink) and reference samples (S1, colored in blue) are overlaid before alignment (top panel), after PASTE alignment (middle panel), or after CAST Stack alignment (bottom panel). **b**, PASTE center-align alignment results of samples S1–S8. Spatial coordinates of these samples are overlaid. Cells are colored by samples. **c**, Manual rotation preprocessing fails to improve PASTE alignment performance. Human-preprocessed spatial coordinates were used as input for PASTE alignment. S7 to S1 alignment is shown as an example. Overlaid spatial coordinates of the two samples are plotted before and after PASTE alignment, the query sample (S7) colored pink and the reference sample (S1) colored blue.

**Extended Data Fig. 6:**
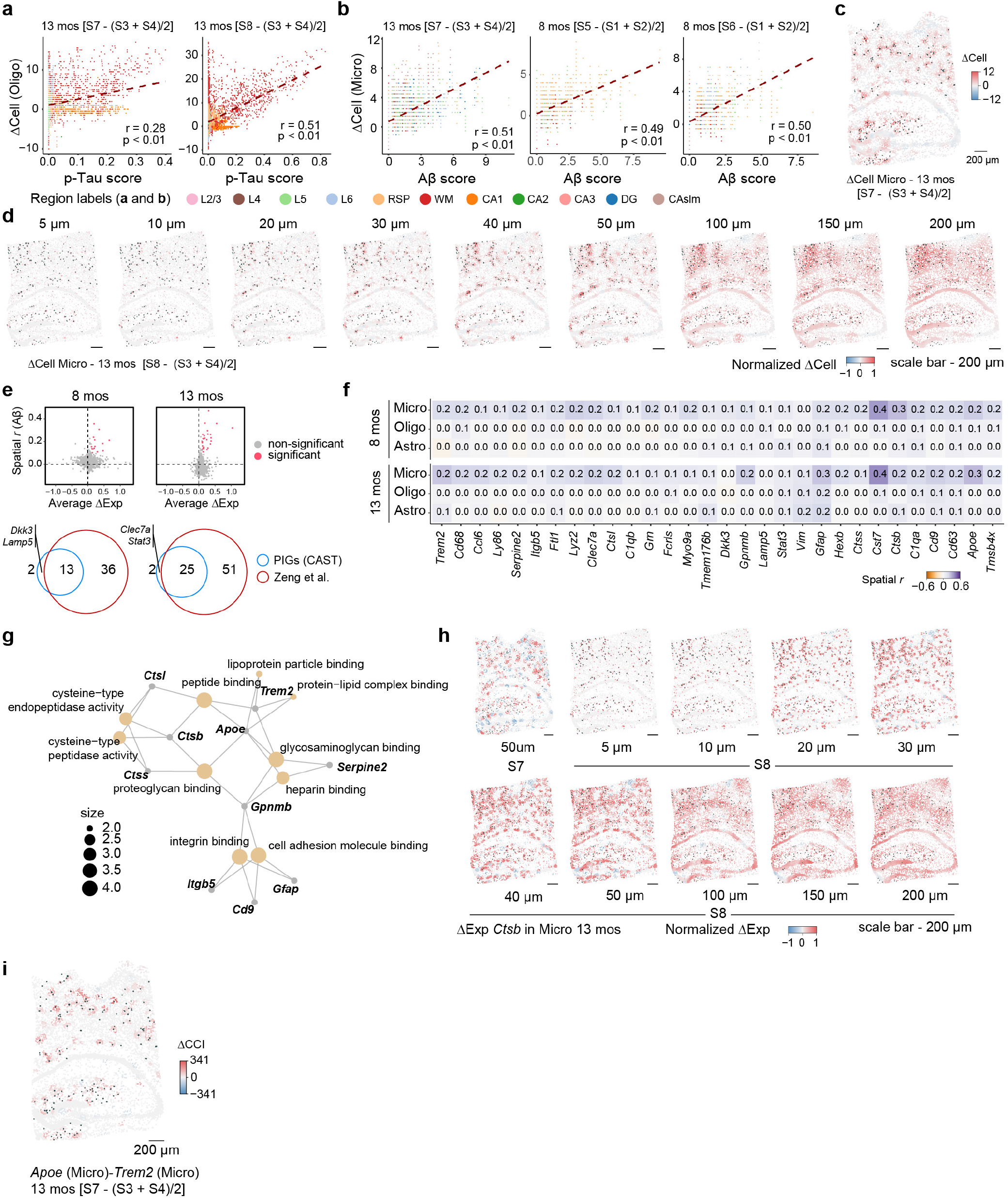
Delta-sample analysis (ΔAnalysis) detects spatial differences of molecular characteristics between disease and normal conditions. **a**, The scatter plot shows the significant high spatial correlation (Pearson correlation test) between the ΔCell (Oligodendrocyte) and the p-Tau score in S7 sample (left panel, n = 9,634) and S8 sample (right panel, n = 10,372). The ΔCell values (*y* axis) are the subtraction of the disease sample and the average value of the control samples (left panel, S7 − (S3 + S4) / 2; right panel, S8 − (S3 + S4) / 2). **b**, Analogous to **a**, the ΔCell (Microglia) shows significant high spatial correlation with Aβ-plaque score in 13 mos (S7 − (S3 + S4) / 2, n = 9,634) and 8 mos (S5 − (S1 + S2) / 2, n = 8,202; S6 − (S1 + S2) / 2, n = 8,186) comparisons. **c**, The spatial gradient map (S7 coordinates) shows the ΔCell pattern of the microglia in 13 mos comparison (S7 − (S3 + S4) / 2). The disease sample shows the microglia abundance in S7, while the control sample shows the average values of the S3 and S4. The dark green dots represent the Aβ-plaque in S8 sample, and the size of the dots indicate the area of the Aβ-plaque. **d**, Analogous to **c**, The ΔCell pattern of the Microglia in 13 mos comparison (S8 − (S3 + S4) / 2) with different physical radii *R* (from 5 μm to 200 μm). **e**, The scatter plots show the overall ΔExp of each gene and its spatial correlation with Aβ plaque score in different comparisons. Venn diagrams show the overlap of the identified plaque-induced genes with the ones identified in initial study. **f**, The average spatial correlation (Pearson *r*) between the ΔExp of each plaque-induced gene and the Aβ-plaque score. For visualization, The values of 4 combinations in each comparison are averaged (13 mos: S7 − S3, S7 − S4, S8 − S3, S8 − S4; 8 mos: S5 − S1, S5 − S2, S6 − S1, S6 − S2). **g**, The GO enrichment analysis of the plaque-induced genes. **h**, Analogous to **c** and **d**, the ΔExp of the *Ctsb* gene in microglia and 13 mos comparison (S7 − (S3 + S4) / 2, *R* = 50 μm; S8 − (S3 + S4) / 2, *R* from 5 μm to 200 μm). **i**, Analogous to **c**, the ΔCCI pattern of the *Apoe* (Micro) - *Trem2* (Micro) in 13 mos comparison (S7 − (S3 + S4) / 2).

**Extended Data Fig. 7:**
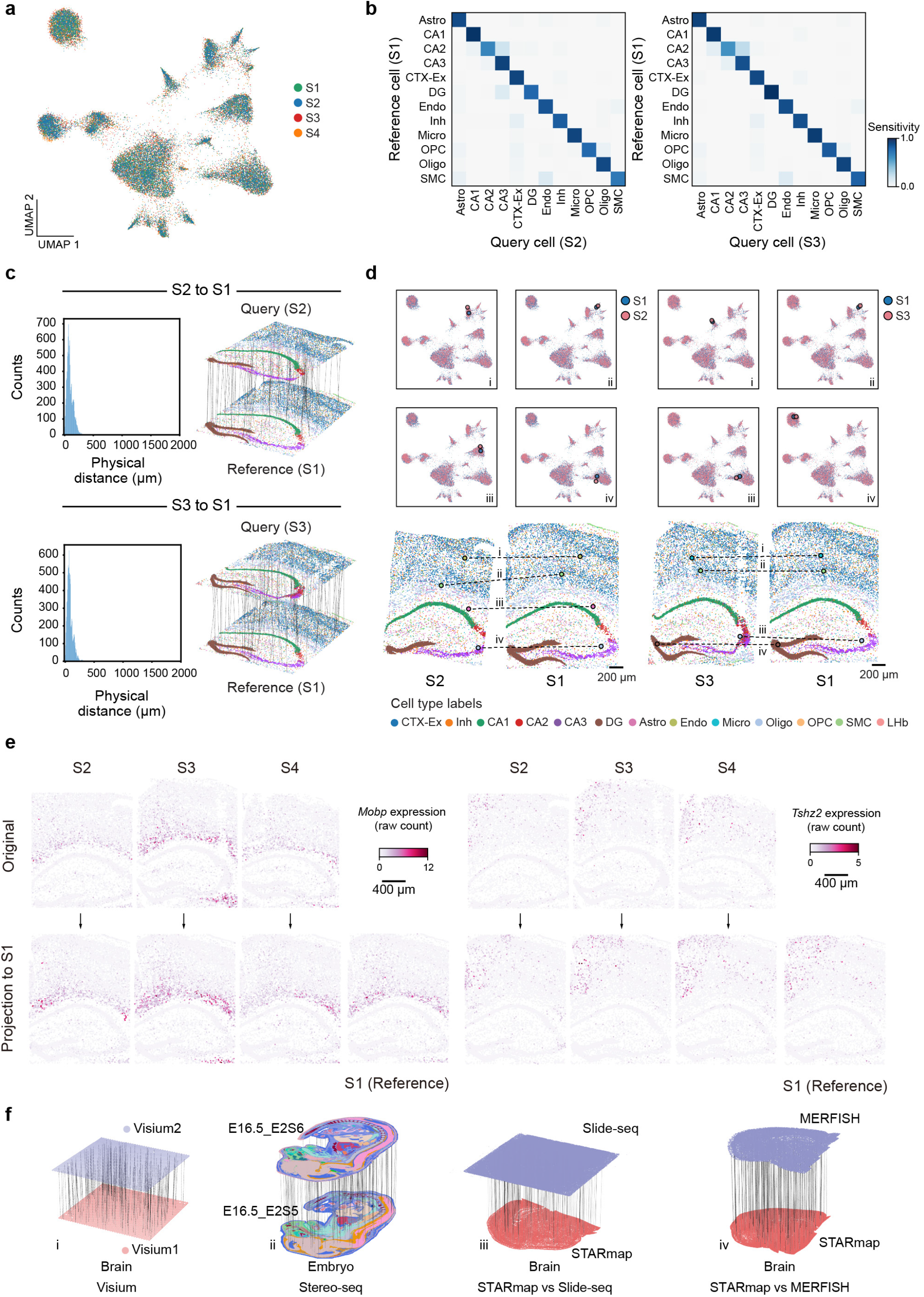
CAST Projection accurately preserves gene expression and spatial and relationships in cells across samples. **a**, UMAP of Combat and Harmony31,32 integrated embedding across S1–S4 samples (control samples) shows that the 4 samples are well integrated in different clusters. Different colors represent the different samples. **b**, Confusion matrix of the projection performance (S2 to S1, True Positive rate (TP) = 0.88; S3 to S1, TP = 0.91). The cell types with more than 10 cells in the reference sample are visualized. **c**, Left panel: The distribution of the physical distance in the spatial single-cell projection of the S2 to S1 (top) and S3 to S1 (bottom). Right panel: Schematic for CAST Projection results. Dashed lines (100 randomly sampled alignment pairs for visualization) connect cells from the query sample (top panel: S2, bottom panel: S3) with its destination cell in the reference sample (S1). **d**, Assignment examples between S2 to S1 (left panel) and S3 to S1 (right panel) in the UMAP plots (top, S2 and S3, light red; S1, blue) and real samples (bottom, the colors of the cells are colored by cell types). **e**, *Mobp* and *Tshz2* gene expression (raw count) profile in the original samples (top, S2–S4) and projected samples (bottom). **f**, CAST Projection is applicable across major spatial technologies and organs. **i**, Visium (Visium1: Mouse Brain Coronal Section 1; Visium2: Mouse Brain Coronal Section 2); **ii**, Stereo-seq; **iii**, STARmap (STARmap_mouse1) vs. Slide-seq; **iv**, STARmap (STARmap_mouse1) vs. MER-FISH. CAST Projection preserves original single-cell resolution on mouse half brain and whole mouse embryo samples. Dashed lines (200 randomly sampled assignment pairs for visualization) connect cells from the query sample with its destination cell in the reference sample.

**Extended Data Fig. 8:**
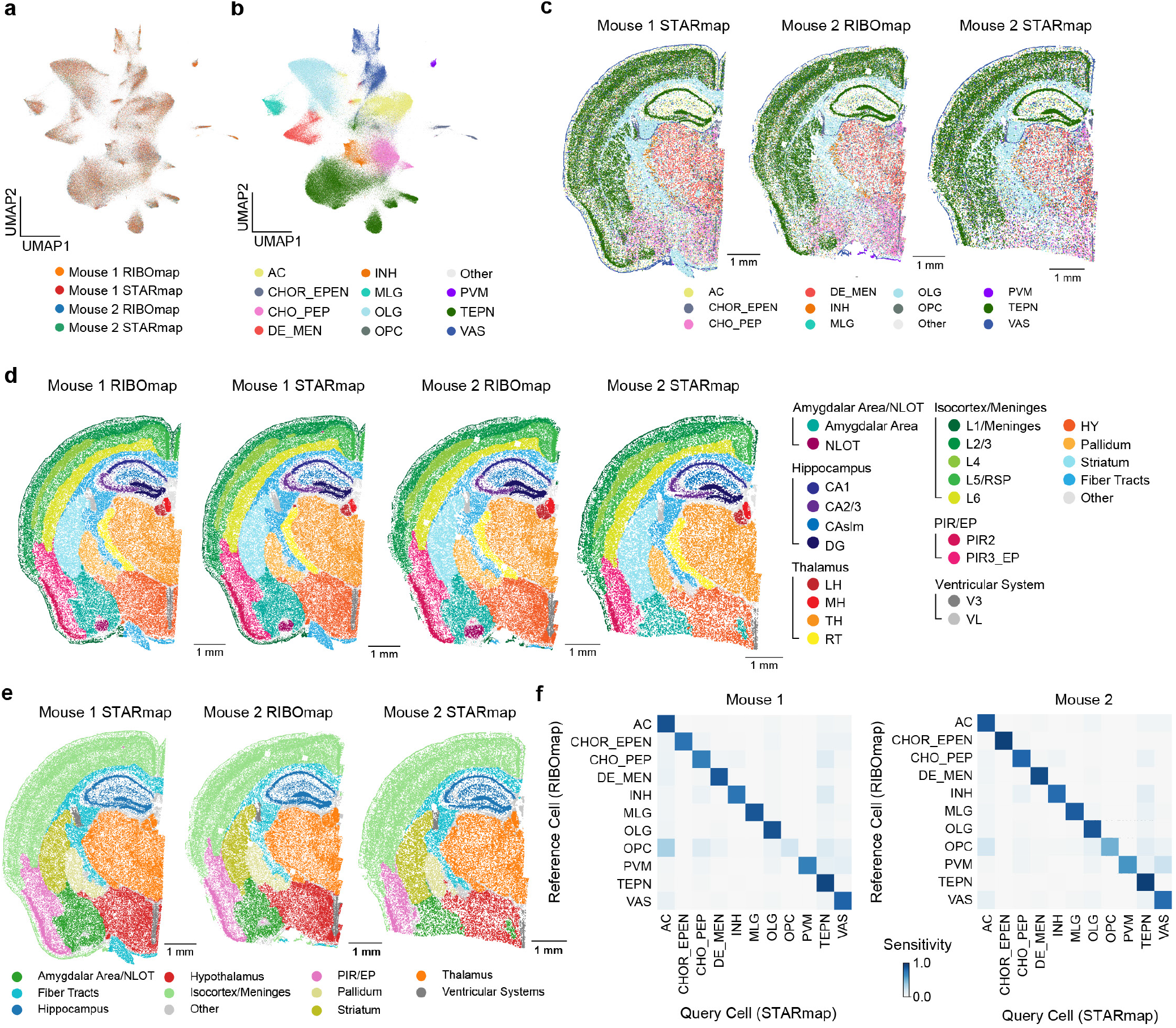
Integration of spatially resolved single-cell ribosome profiling and gene expression profiling. **a**,**b**, UMAP visualization of integrated RIBOmap and STARmap datasets. Each cell is colored by the dataset (**a**) or cell type (**b**). **c**, Cell type profiles of Mouse1 STARmap, Mouse2 RIBOmap and Mouse2 STARmap samples. **d**,**e**, Tissue region (**e**) and sub-region (**d**) profiles of all 4 samples generated using CAST Mark. The hierarchy of tissue regions are shown in the legends (right panel, **d**). HY, Hypothalamus; LH, Lateral habenula; MH, Medial habenula; NLOT, Nucleus of the lateral olfactory tract; PIR2, Piriform area, pyramidal layer; PIR3_EP, Piriform area, polymorph layer and Endopiriform nucleus; RT, Reticular nucleus of the thalamus; TH, Thalamus; V3, third ventricle; VL, lateral ventricle. **f**, Confusion matrix of the projection results (Mouse1, True Positive rate (TP) = 0.84; Mouse2, TP = 0.86).

**Extended Data Fig. 9:**
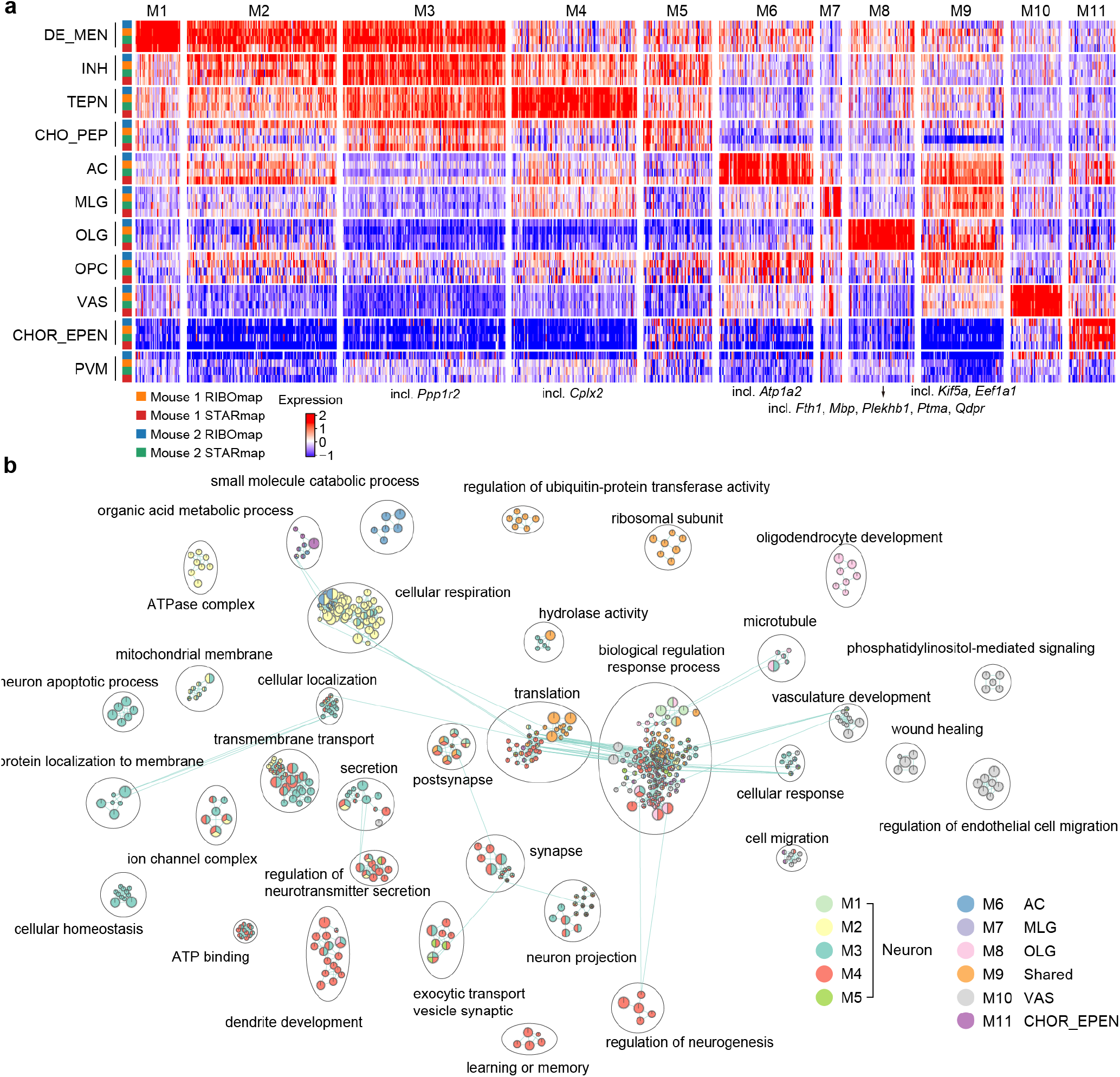
Gene modules identified by co-clustering of RIBOmap and STARmap data in four samples. **a**, Genes are clustered into 11 distinct clusters by co-clustering of RIBOmap and STARmap data in four samples (Z-score expression). Example genes shown in Fig. 6b are marked below the gene modules they belong to. **b**, Enriched GO terms in each gene module are grouped by terms and colored by gene modules. In the enrichment map, nodes represent the enriched GO terms, while the size of the node corresponds to the number of genes in the GO terms. The edges between nodes indicate the overlapping genes between the GO terms.

**Extended Data Fig. 10:**
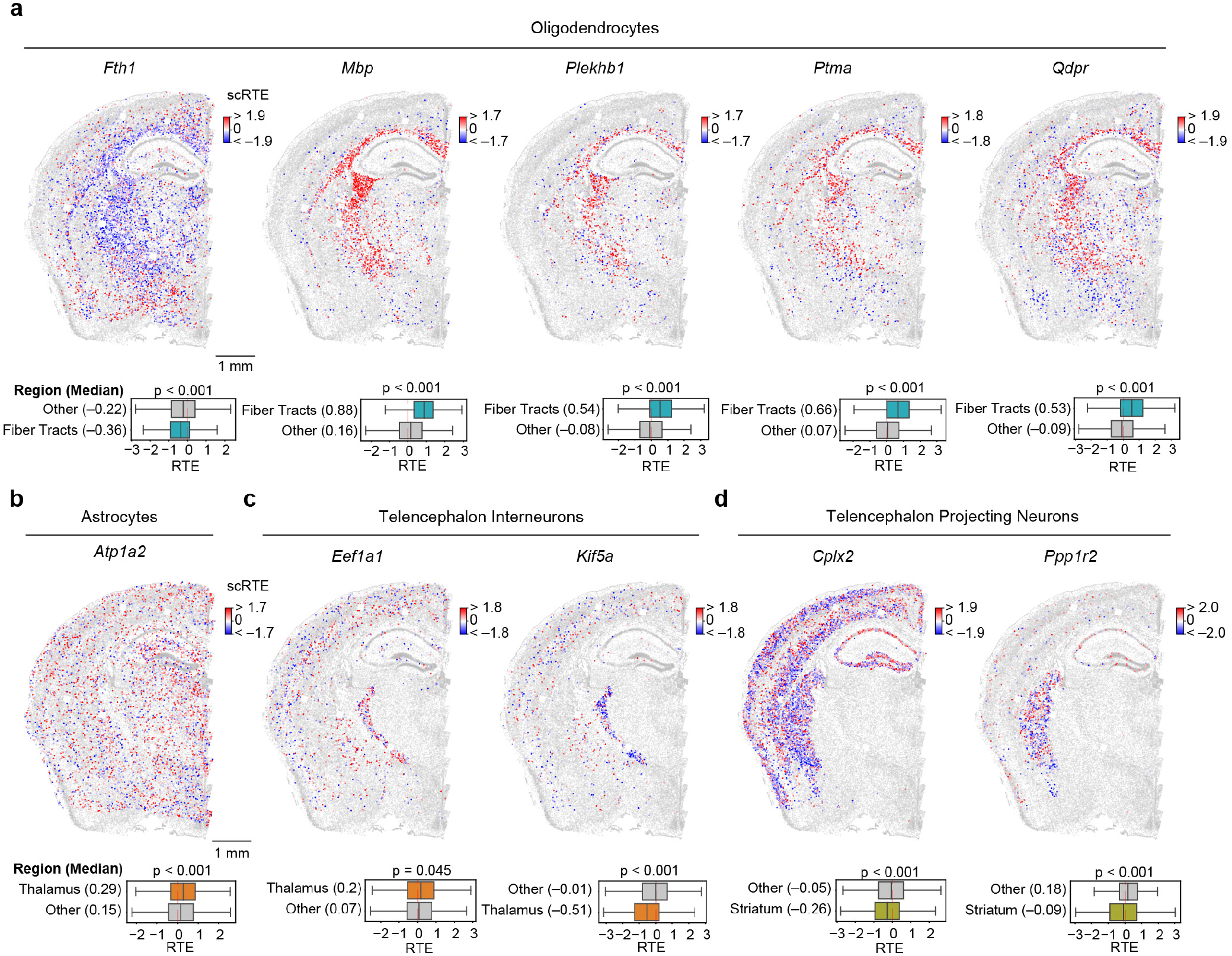
Examples of the cell-type specific scRTE patterns across different genes. **a**-**d**, Spatially resolved and cell type specific scRTE profiles in mouse2. Cells of the annotated cell type (above) and with available scRTE values are colored by scRTE levels, other cells are colored gray. The boxplots with Kruskal-Wallis test are used to evaluate the differences across the groups.

**Supplementary Table 1.**
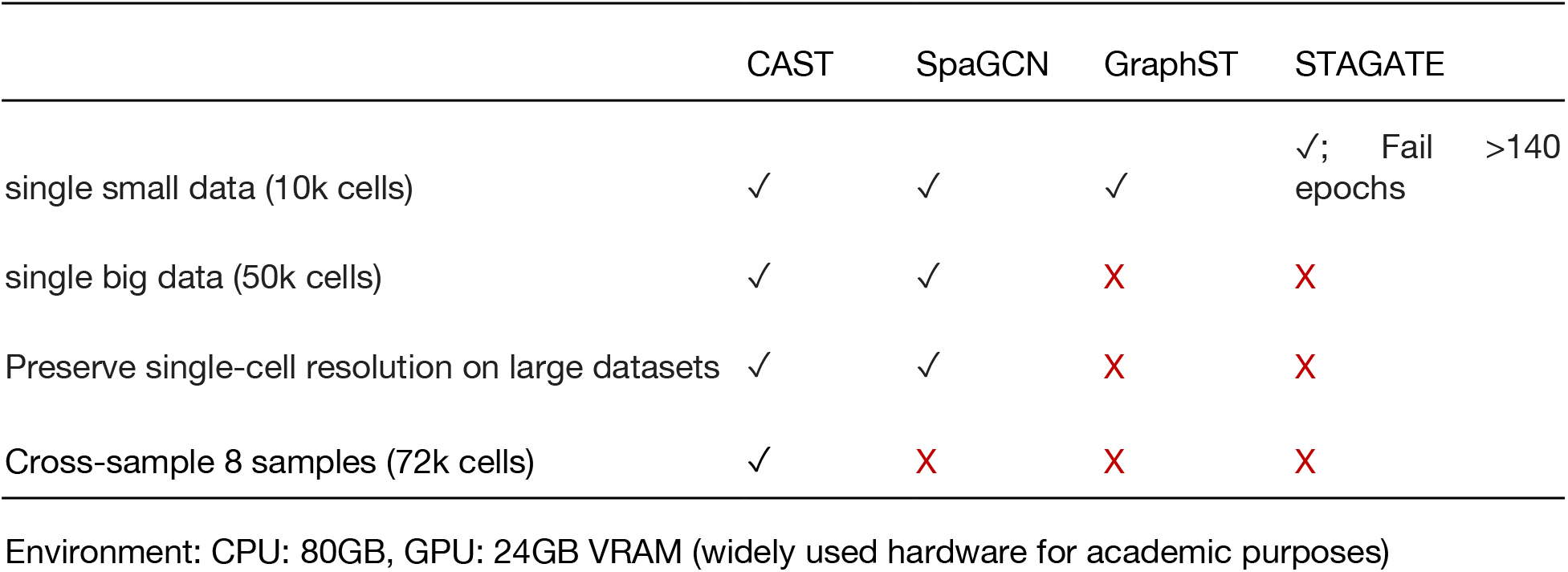
Tissue segmentation benchmark.

**Supplementary Table 4.**
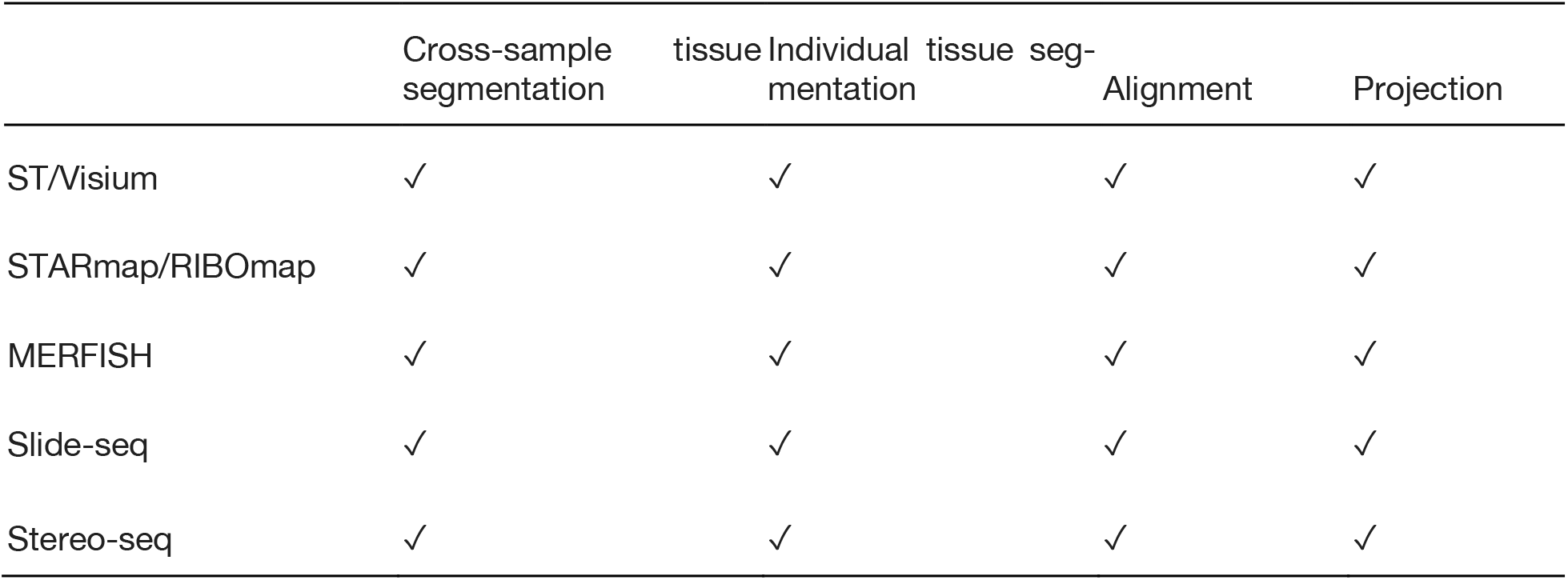
Broad applicability of CAST.

**Supplementary Table 5.**
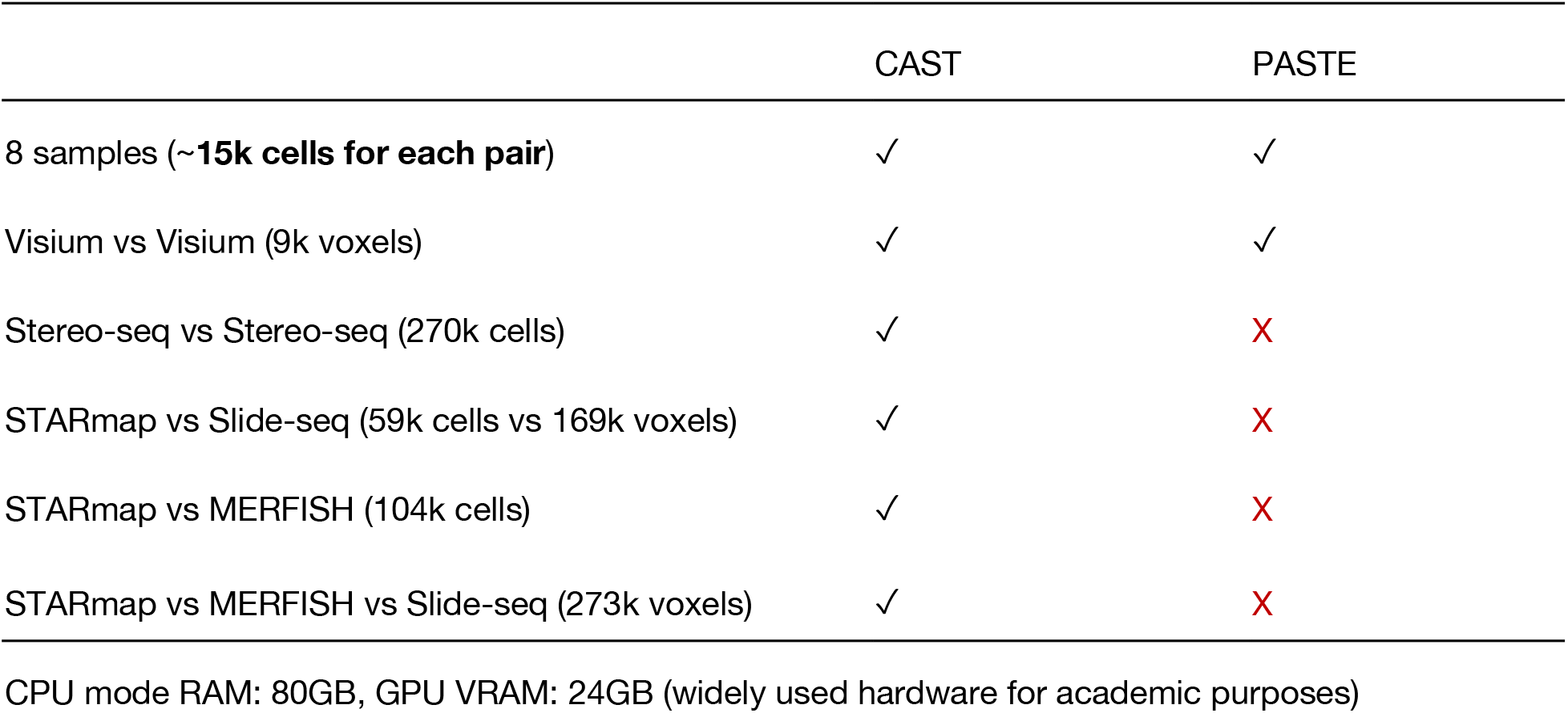
Sample alignment benchmark.

|  | CAST | PASTE |
| --- | --- | --- |
| 8 samples (~**15k cells for each pair**) | ✓ | ✓ |
| Visium vs Visium (9k voxels) | ✓ | ✓ |
| Stereo-seq vs Stereo-seq (270k cells) | ✓ | X |
| STARmap vs Slide-seq (59k cells vs 169k voxels) | ✓ | X |
| STARmap vs MERFISH (104k cells) | ✓ | X |
| STARmap vs MERFISH vs Slide-seq (273k voxels) | ✓ | X |

CPU mode RAM: 80GB, GPU VRAM: 24GB (widely used hardware for academic purposes)

